# Controllable self-replicating RNA vaccine delivered intradermally elicits predominantly cellular immunity

**DOI:** 10.1101/2022.09.05.506686

**Authors:** Tomokazu Amano, Hong Yu, Misa Amano, Erica Leyder, Maria Badiola, Priyanka Ray, Jiyoung Kim, Akihiro C. Ko, Achouak Achour, Nan-ping Weng, Efrat Kochba, Yotam Levin, Minoru S.H. Ko

**Affiliations:** Elixirgen Therapeutics, Inc., Baltimore, MD, USA; Laboratory of Molecular Biology and Immunology, National Institute on Aging, NIH, Baltimore, MD, USA; NanoPass Technologies Ltd., NanoPass Technologies Ltd, Nes Ziona, Israel

**Keywords:** Vaccine, Self-replicating RNA, Temperature-sensitive, Temperature-controllable, COVID-19 Coronavirus, Intradermal, Skin, T-cell-inducing, Cellular Immunity, pan-coronavirus vaccine

## Abstract

Intradermal delivery of self-replicating RNA (srRNA) is a promising vaccine platform. Considering that human skin temperature is around 33°C, lower than core body temperature of 37°C, we have developed an srRNA that functions optimally at skin temperature and is inactivated at or above 37°C as a safety switch. This temperature-<u>c</u>ontrollable srRNA (c-srRNA), when tested as an intradermal vaccine against SARS-CoV-2, functions when injected naked without lipid nanoparticles. Unlike most currently available vaccines, c-srRNA vaccines predominantly elicit cellular immunity with little or no antibody production. Interestingly, c-srRNA-vaccinated mice produced antigen-specific antibodies upon subsequent stimulation with antigen protein. Antigen-specific antibodies were also produced when B-cell stimulation using antigen protein was followed by c-srRNA booster vaccination. Using c-srRNA, we have designed a pan-coronavirus booster vaccine that incorporates both spike receptor binding domains as viral surface proteins and evolutionarily conserved nucleoproteins as viral non-surface proteins, from both SARS-CoV-2 and MERS-CoV. It can thereby potentially immunize against SARS-CoV-2, SARS-CoV, MERS-CoV, and their variants. c-srRNA may provide a route to activate cellular immunity against a wide variety of pathogens.

## Introduction

Betacoronaviruses encompass SARS-CoV-2; SARS-CoV, which caused the 2002–2004 SARS outbreak; and Middle East respiratory syndrome (MERS)-CoV, which has occasionally caused fatal illness (Gorbalenya et al., 2020). The current pandemic stimulated the application of existing vaccine platforms towards SARS-CoV-2 as well as the development of new types of vaccines – most notably, the mRNA-based vaccines developed by Pfizer/BioNTech (Walsh et al., 2020) and Moderna (Jackson et al., 2020) were among the first introduced and widely used (Hogan and Pardi, 2021) (Szabó et al., 2022). As an alternative RNA vaccine technology, self-replicating RNA (srRNA) or self-amplifying RNA (saRNA or SAM) has also been used (de Alwis et al., 2021)(Mckay et al., 2020). Unlike mRNA, srRNA encodes both a gene(s) of interest and an RNA-dependent RNA polymerase, and thus provides additional features such as longer expression and no use of modified nucleotides (reviewed in (Pushko and Tretyakova, 2014)(Brito et al., 2015)(Lundstrom, 2016)(Ballesteros-Briones et al., 2020)(Blakney et al., 2021).

Although the current mRNA vaccines are highly potent inducers of neutralizing antibody (NAb) production, the immunity wanes significantly over a period of months, and they only weakly induce cellular immunity (Levin et al., 2021; Nathan et al., 2021). But the critical importance of cellular immunity in fighting coronaviruses has been demonstrated experimentally and extensively discussed (Sette and Crotty, 2021)(Moss, 2022). Cellular immunity depends on linear T-cell epitopes, whereas B-cell humoral immunity also responds to conformational epitopes. Consequently, cellular immunity is much more robust against variant viruses than is humoral immunity. In addition, cellular immunity alone can provide protection via the CD8+ cytotoxic T-cell-mediated destruction of infected cells (Channappanavar et al., 2014)(Matchett et al., 2021). Furthermore, memory T cells last longer than memory B cells, potentially providing lifelong immunity (Channappanavar et al., 2014). Therefore, the development of vaccines that induce strong cellular immunity, “T-cell-inducing vaccines”, has been one of the major goals for vaccine developers (Gilbert, 2012).

One way to achieve such cellular immunity is through intradermal vaccination (Hickling et al., 2011)(Dugan et al., 2020). Human skin (epidermis and dermis) is rich in antigen-presenting cells (APCs), including Langerhans cells and dermal dendritic cells; about 40% of the body’s APCs are in the skin (Chen and Wu, 2011). Because it targets the APCs, intradermal vaccination is considered more effective than subcutaneous or intramuscular vaccination (Chen and Wu, 2011; Hung and Yuen, 2018)(Hickling et al., 2011), and such targeting activates the T-cell immunity pathway. In addition, modern intradermal injection devices such as the MicronJet600 (Levin et al., 2015) and Immucise (Shimizu et al., 2022) lower the skill level required for injection compared to the traditional Mantoux method.

In contrast to the standard method to enhance incorporation of mRNAs into cells by formulating them with a lipid nanoparticle, intradermal injection using the MicronJet600 makes naked mRNAs (no lipid nanoparticles, no adjuvant) as immunogenic as mRNAs formulated with lipid nanoparticles (Golombek et al., 2018). Similarly, intradermal injection of naked srRNA followed by electroporation elicits strong immunogenicity (Johansson et al., 2012). In fact, LNP-formulated srRNA shows only several-fold higher expression than naked srRNA (Blakney et al., 2019). mRNA or srRNA formulated without LNP or adjuvants could thus potentially be as effective while reducing safety risks and skin irritation. After intradermal administration of srRNA, the srRNA is predominantly taken up by immune cells including the skin APCs (Blakney et al., 2019; Leyman et al., 2018).

We have added one feature to srRNAs that had not been previously considered. Unlike the core body temperature of 37°C, the temperature at the surface of the body varies in different areas of skin, in different conditions (e.g., after exercise), in people with different amounts of body fat, or at different ambient temperatures, but falls within the range of 30°C-35°C (Costa et al., 2018). As far as we know, no vaccine or medication has taken this low skin/body surface temperature into consideration. Here, we have developed an srRNA vector that is optimized to function at 30°C-35°C.

This temperature-<u>c</u>ontrollable self-replicating RNA is called c-srRNA. We have developed several vaccines against SARS-CoV-2 using c-srRNA as a platform, and show that intradermally administered c-srRNA indeed possesses the desirable features of a T-cell-inducing vaccine. Finally, in line with the current shift of global vaccine efforts to the development of so called “pan-coronavirus vaccines,” which can potentially address all major betacoronaviruses and their variants (Dolgin, 2022), we have designed a pan-coronavirus, T-cell inducing booster vaccine based on the c-srRNA vaccine platform.

## Results

### Optimization of self-replicating RNA vector for intradermal delivery

Most self-replicating RNA vectors are based on a Venezuelan Equine Encephalitis virus (VEEV) backbone, in which a subgenomic region encoding structural proteins is replaced with a gene of interest (reviewed in (Pushko and Tretyakova, 2014)(Brito et al., 2015)(Lundstrom, 2016)(Blakney et al., 2021)). We made our initial RNA vector based on the commonly used TRD strain (Kinney et al., 1989) with slight sequence modifications, A551D and P1308S, that are also found in other srRNA vectors (Yoshioka et al., 2013)(Mckay et al., 2020), and a truncated 3’-UTR that is shorter than that in typical srRNA vectors. To produce an srRNA that functions optimally at about 33°C (skin temperature), we systematically mutated the non-structural proteins of our initial RNA vector and tested the expression levels at 33°C and 37°C of the gene of interest encoded in the sub-genomic region. As a guide, we used a published database of a total of 7,480 mutants that were produced by VEEV viral replication at 30°C or 40°C (Data Set S1 from (Beitzel et al., 2010). We identified a mutant with a 15 bp (5 a.a.) insertion in the nsP2 protein that causes the desired temperature sensitivity: it functions at 30-35°C but is inactivated at ≥37°C (Supplemental FIG. S1). We called this *in vitro* transcribed mRNA a controllable srRNA (c-srRNA), more specifically c-srRNA1 (Table 1).

**Table 1.** srRNAs and c-srRNAs developed and used in this study.

| Name | Vector | Gene of interests or antigens |
| --- | --- | --- |
| c-srRNA1-LUC | c-srRNA1 | LUC |
| c-srRNA3-LUC | c-srRNA3 | LUC |
| c-srRNA1-GFP | c-srRNA1 | GFP |
| T7-VEE-GFP-ts | T7-VEE-GFP-ts | GFP |
| c-srRNA1-RBD (EXG-5003) | c-srRNA1 | RBD |
| srRNA0-RBD | srRNA0 | RBD |
| c-srRNA3-RBD | c-srRNA3 | RBD |
| EXG-5003o | c-srRNA3 | RBD (omicron) |
| EXG-5004 | c-srRNA3 | SARS2-N (no CD5sp) |
| EXG-5005 | c-srRNA3 | SARS2-N |
| EXG-5006a | c-srRNA3 | SARS2-N, MERS-N |
| EXG-5006b | c-srRNA3 | SARS2-N, MERS-N (codon optimized) |
| EXG-5008 | c-srRNA3 | SARS2-RBD, SARS2-N, MERS-N, MERS-RBD |

Notes.
LUC, luciferase; GFP, Green Fluorescent Protein; RBD, Spike Receptor Binding Domain of SARS-CoV-2; SARS2-N, nucleoprotein of SARS-COV-2; MERS-N, nucleoprotein of MERS-CoV; SARS2-RBD, Spike Receptor Binding Domain of SARS-CoV-2 (also noted just RBD); MERS-RBD, Spike Receptor Domain of MERS-CoV. CD5sp, Signal peptide of human CD5 protein.

To test whether c-srRNA1 functions *in vivo* in skin, we administered c-srRNA1 encoding the luciferase gene (called c-srRNA1-LUC) intradermally into mice. We formulated c-srRNA1-LUC in naked RNA form in Lactated Ringer’s solution, without LNP or any transfection reagents. [Lactated Ringer’s solution was selected based on previous reports (Probst et al., 2007) (Phua et al., 2013).] As a control, 5MoU-modified synthetic mRNA encoding LUC was used (called mRNA-LUC). 5 μg of mRNAs encoding luciferase, either c-srRNA1-LUC or mRNA-LUC, were injected intradermally at a single site on the right hind limb of CD-1 outbred mice. Luciferase activity was visualized and quantitated using a bioluminescent imaging system (AMI HTX). Luciferase imaging showed that intradermal injection of naked RNA encoding luciferase resulted in expression of luciferase *in vivo* (FIG. 1). Strikingly, the *in vivo* expression of luciferase driven by c-srRNA1-LUC continued for nearly a month, apparently sustained by its self-replication. In contrast, the *in vivo* expression of LUC driven by mRNA-LUC lasted only a week. Furthermore, the expression level of LUC in recipients of c-srRNA1-LUC was 10− to 100− fold higher than the expression level of luciferase in recipients of mRNA-LUC. Importantly, luciferase expression was not observed in the un-injected areas of the recipients’ skin nor in other regions of mice, including the internal organs (FIG. 1). Furthermore, direct injection into skeletal muscle also did not show luciferase activities (Supplemental FIG. S2). This observation indicates that, as expected, the temperature-sensitive c-srRNA1-LUC did not replicate and express luciferase in non-permissive conditions.

**FIG. 1.**
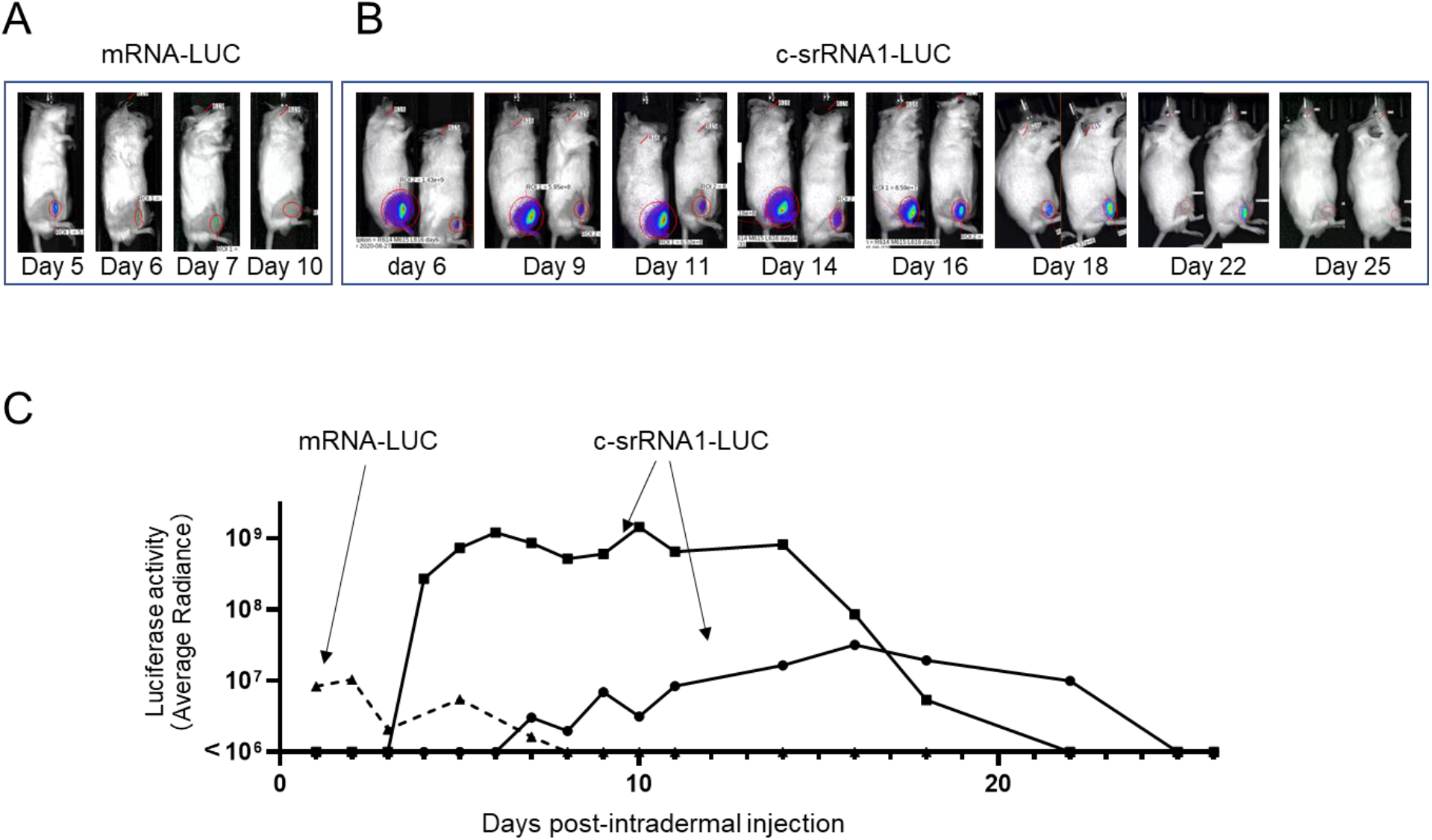
In vivo expression of LUC after intradermal injection of c-srRNA. mRNAs, either mRNA-LUC (CleanCap Fluc mRNA [5MoU], TriLink L-7202) or c-srRNA1-LUC, were formulated as naked RNAs, without LNP or other transfection reagents, in Lactated Ringer’s solution. 5 μg of mRNAs were injected intradermally into a single site on the right hind limb of CD-1 outbred mice (Day 0). Luciferase activity was visualized and quantitated by using a bioluminescent imaging system, AMI HTX (Spectral Instruments Imaging, Tucson, AZ) from Day 1 through Day 26. (**A**) Pictures of mice that received intradermal injection of mRNA-LUC. (**B**) Pictures of mice that received intradermal injection of c-srRNA1-LUC. (**C**) Changes of LUC activities over time. Luciferase activity was represented as average radiance.

### Generation of c-srRNA1 vaccine encoding SARS-CoV-2 spike receptor binding domain

To test the suitability of c-srRNA1 vectors for vaccination, we generated a c-srRNA1 encoding the receptor binding domain (RBD) of the spike protein of severe acute respiratory syndrome coronavirus-2 (SARS-CoV-2, also known as 2019-nCoV). At the time of antigen selection, no report on a vaccine against SARS-CoV-2 was published, but we considered the RBD as the most reasonable vaccine antigen (see the Materials and Methods section for the detailed line of thinking). We used the originally reported Wuhan strain sequence (NCBI accession number: NC_045512, (Wu et al., 2020)). To make the antigen as a secreted protein, the signal peptide sequence of the human CD5 protein (NCBI accession number: NM_014207.4) was fused to the N-terminus of the RBD. This c-srRNA1-RBD is named EXG-5003 (Table 1).

### EXG-5003 induces cellular immunity

Because our initial goal was to develop a T-cell-inducing vaccine, we tested whether the intradermal injection of EXG-5003 RNA could induce cellular immunity (FIG. 2A).

**FIG. 2.**
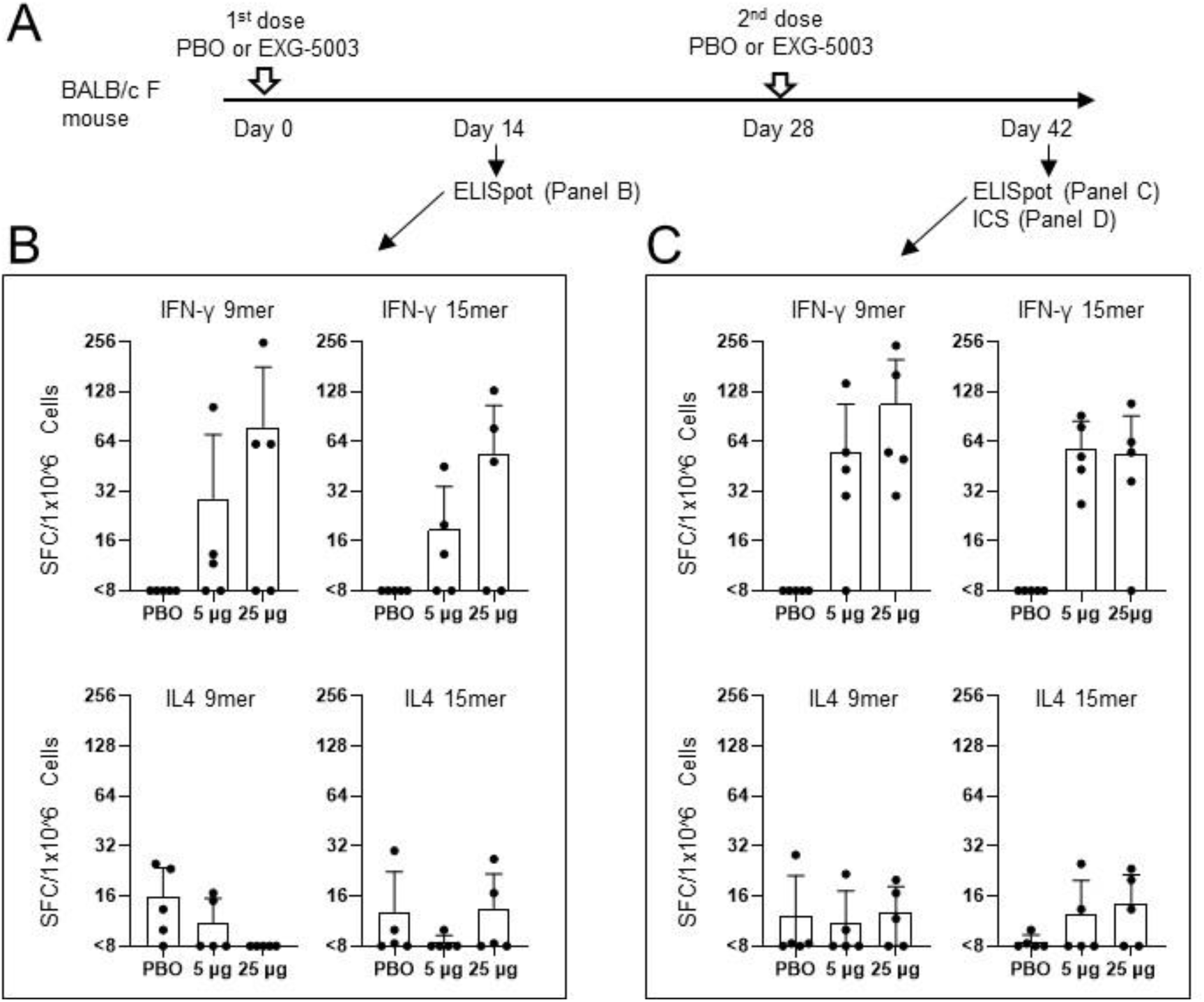
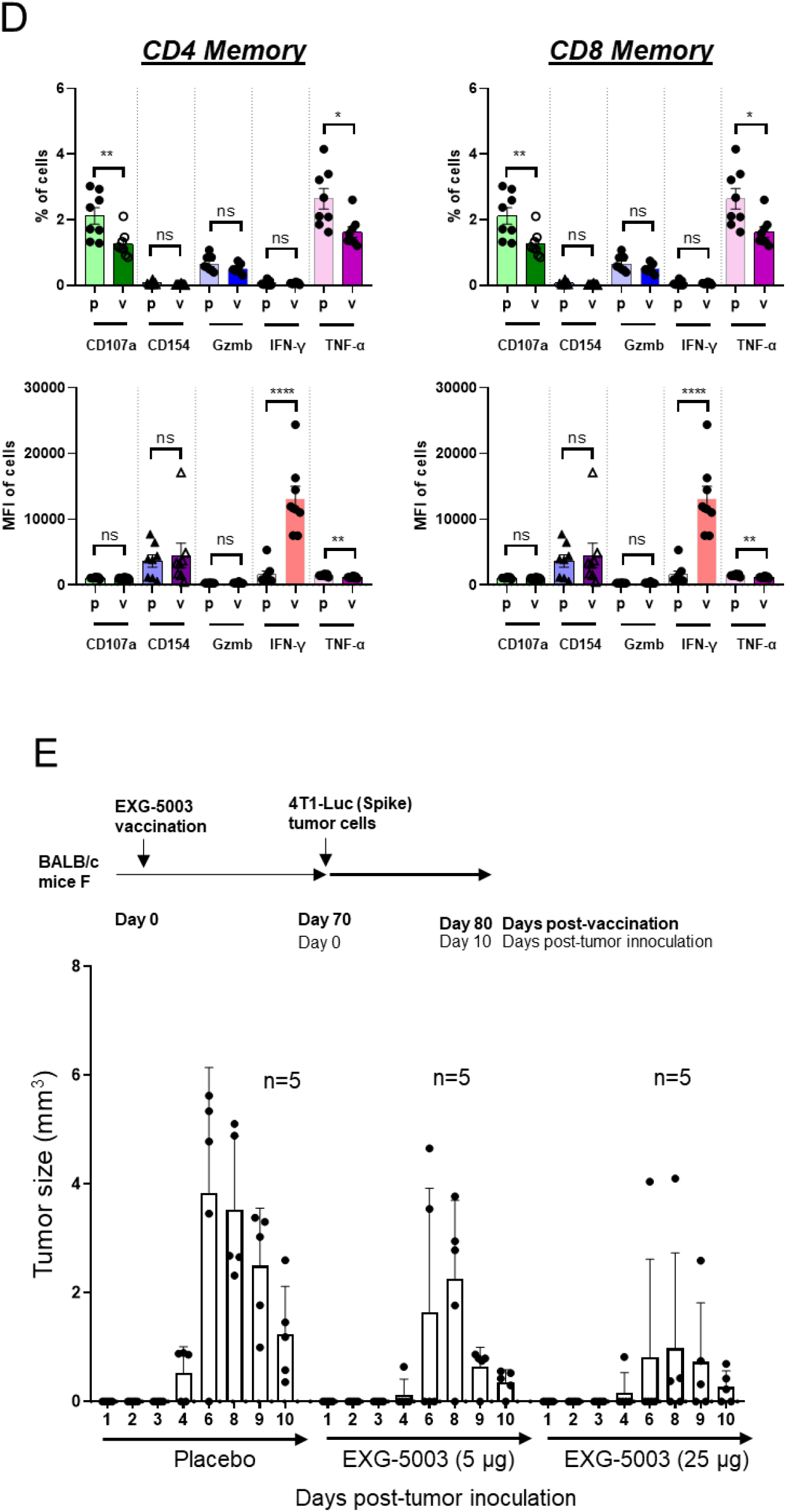
c-srRNA vaccine elicits cellular immunity. (**A, B, C**) Cellular immunity assessed by enzyme-linked immunospot (ELISpot) assays. A single dose or two doses of a placebo (PBO) or 5 μg, or 25 μg of EXG-5003 RNAs, formulated as naked RNAs without LNP or other transfection reagents and in Lactated Ringer’s solution, were administered intradermally to BALB/c mice. To assess cellular immunity, an ELISPOT assay, which quantitates the number of cytokine secreting cells, was performed on splenocytes isolated on Day 14 and Day 42. The splenocytes were stimulated for 24 hours with pools of peptides: 9mer peptides for RBD of SARS-CoV-2 (original strain) or 15mer peptides for RBD of SARS-CoV-2 (original strain). (**D**) Cellular immunity was assessed by intracellular cytokine staining and flow cytometry (ICS) assays. The FACS-ICS assays were performed on splenocytes isolated on Day 42. MFI, mean fluorescence intensity; p, placebo; v, EXG-5003. (**E**) Cellular immunity assessed by *in vivo* tumor cell elimination assays. A single EXG-5003 vaccination suppresses the growth of 4T1-LUC tumor cells carrying SARS-CoV-2 spike protein (4T1-LUC-Spike cells).

First, we assessed the induction of cellular immunity by the enzyme-linked immunospot (ELISpot) assay. T-cell epitopes are short linear peptides with size ranges of 8mer - 11mer for MHC-I and 10mer – 30mer for MHC-II; therefore, we tested both 9mer and 15mer peptide libraries to restimulate the T cells in culture. As shown in FIG.2B and 2C, EXG-5003 RNA administered by intradermal injection induced cellular immunity against the SARS-CoV-2 RBD. IFN-γ-secreting cells, which are characteristic of CD8+ cytotoxic T cells and type 1 CD4+ helper T cells (Th1 cells), were induced by the EXG-5003 RNA vaccine. In contrast, IL4-secreting cells, which are characteristic of type 2 CD4+ T helper cells (Th2 cells), were only weakly induced by the EXG-5003 vaccine. The results showed that intradermal administration of EXG-5003 elicited a Th1 dominant (Th1>Th2) cellular immune response against SARS-CoV-2 RBD, a favorable feature for a vaccine directed against a viral pathogen.

Next, we examined the induction of cellular immunity by Intracellular Cytokine Staining and Flow Cytometry (ICS) assays. Both CD4+ T cells and CD8+ T cells showed an antigen-specific increase of mean fluorescence intensity (MFI) of IFN-γ secretion by day 14 of the 2^nd^ dosing (FIG. 2D). As expected at this early time point post-vaccination, TNF-α did not show such antigen-specific increase (e.g., (Rodrigues et al., 2021)).

We then assessed whether EXG-5003 can induce RBD-specific CD8+ cytotoxic T cells, which can eliminate cells infected with SARS-CoV-2. As an experimental model, we used 4T1 mouse mammary tumor cells that were transfected with a plasmid vector expressing SARS-CoV-2 spike protein (called 4T1-LUC-Spike cells). Mice were first vaccinated with EXG-5003. Seventy days later, 4T1-LUC-Spike cells were injected into mice and tumor growth assessed by measuring tumor size. A single dose of EXG-5003 was sufficient to suppress the subsequent growth of 4T1-LUC-Spike tumor cells carrying SARS-CoV-2 spike protein (FIG. 2E). We inferred that the intradermal injection of EXG-5003 induced RBD-specific CD8+ cytotoxic T cells.

### EXG-5003 does not induce but primes humoral immunity

We also assessed whether EXG-5003 can induce humoral immunity, i.e., RBD-specific antibodies. As shown in FIG. 3, no increase of RBD-specific IgG was observed in any group treated with two vaccine doses of EXG-5003 (FIG. 3; Day −3, Day 14, Day 28, Day 46). We also tested whether the addition of RNase inhibitor can enhance immunogenicity, as it has been previously shown that an RNase-inhibitor can enhance the *in vivo* stability of srRNA in mouse skin (Huysmans et al., 2019). Again, no RBD-specific IgG was seen. Thus, intradermal administration of EXG-5003 RNA does not induce humoral immunity.

**FIG. 3.**
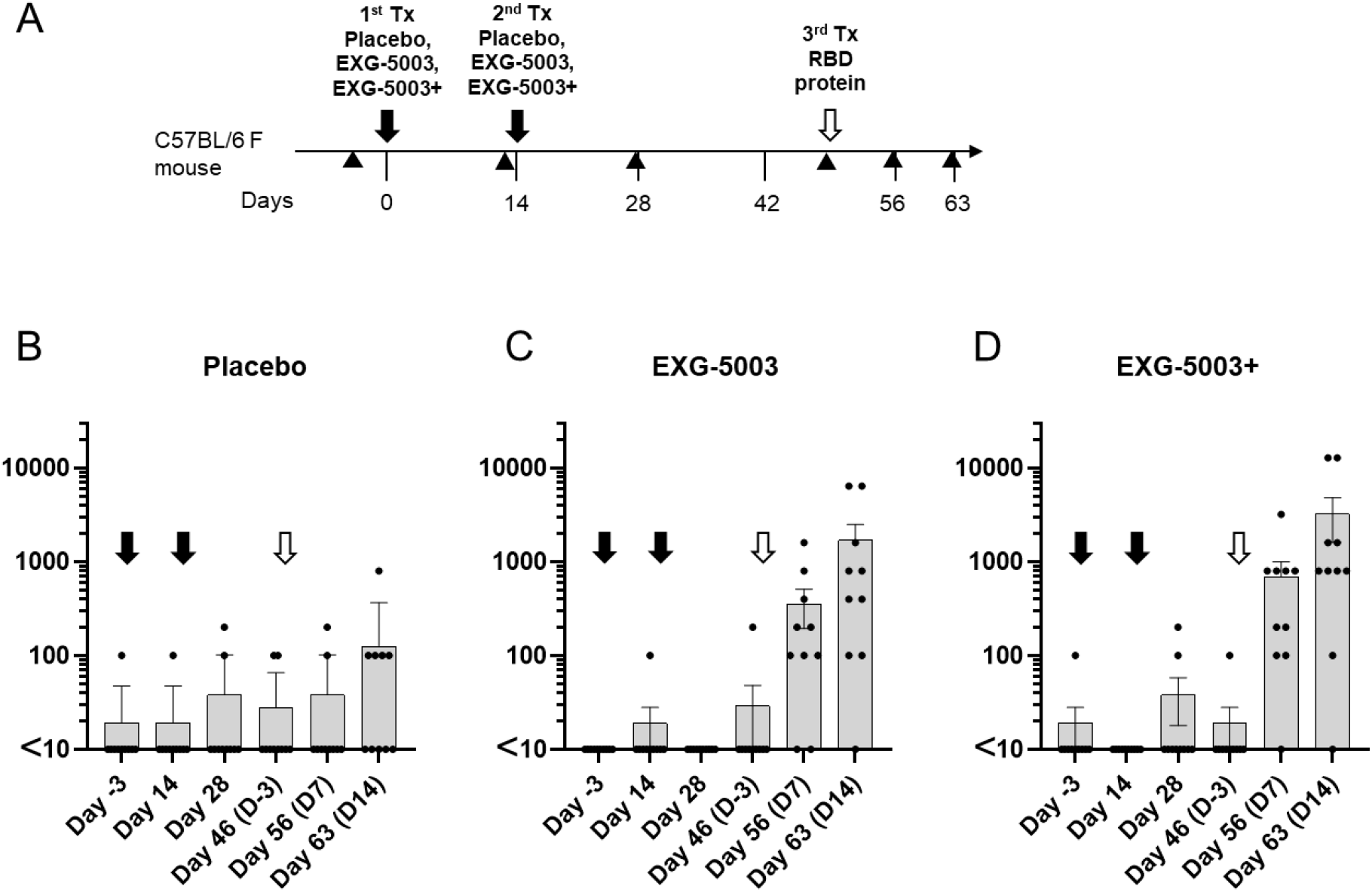
c-srRNA vaccines prime humoral immunity, thereby inducing antigen-specific antibody when further triggered by the antigen protein. (**A**) A schematic diagram of experimental procedures. EXG-5003 RNAs were formulated as naked RNAs, without LNP or other transfection reagents, in Lactated Ringer’s solution. (**B**) Groups of CD-1 outbred mice (N = 10) received one of three formulations by intradermal injection on Day 0 and Day 14 (black arrows): a placebo (buffer only), 5 μg of EXG-5003 RNA, or 5 μg of EXG-5003 RNA in combination with an RNase inhibitor (3 units of RNasin Plus; Promega, Madison, WI). Subsequently, all mice received a recombinant RBD protein (Ala319-Phe541, with a C-terminal 6-His tag, Accession # YP_009724390.1: R&D Systems, Minneapolis, MN) by intradermal injection on Day 49 (open arrows). To assess humoral immunity, an enzyme-linked immunosorbent assay (ELISA), which quantitates the amount of immunoglobulin G (IgG) specific to a recombinant RBD protein (represented as the geometric mean of endpoint titer in triplicate), was performed on serum obtained from the mice on Day −3, Day 14, Day 28, Day 46, Day 56, and Day 63.

In contrast to its inability to induce humoral immunity on its own, when mice that received prior intradermal injections of EXG-5003 then received a booster comprising a recombinant RBD protein (on Day 49), they quickly responded with an increase in RBD-specific IgG (FIG. 3C, D), whereas the placebo group showed only a slight increase in RBD-specific IgG (FIG. 3B). In this case, addition of RNase inhibitor indeed provided mild enhancement of EXG-5003 RNA function (FIG. 3D). We infer that the mice that received EXG-5003 vaccinations maintained immune memory and can thus exhibit a secondary humoral response to the recombinant RBD protein.

### EXG-5003 primes humoral immunity against delta variant spike protein

Prompted by the unequivocal finding that intradermal c-srRNA vaccination primes humoral immunity, we checked whether post-vaccination exposure to a variant antigen slightly different from the antigen encoded on the vaccine can induce antibodies against the variant antigen. BALB/c mice were first treated with intradermal injection of EXG-5003 RNA, which encoded the RBD of SARS-CoV-2 [original strain] and was formulated as a naked RNA without LNP nor transfection reagent. The mice were then treated with the variant RBD protein of SARS-CoV-2 (delta variant B.1.617.2) along with adjuvant (FIG. 4A). As expected, cellular immunity assessed by the presence of antigen-specific IFN-γ-secreting T cells was already induced by day 14 post-vaccination (FIG. 4B).

**FIG. 4.**
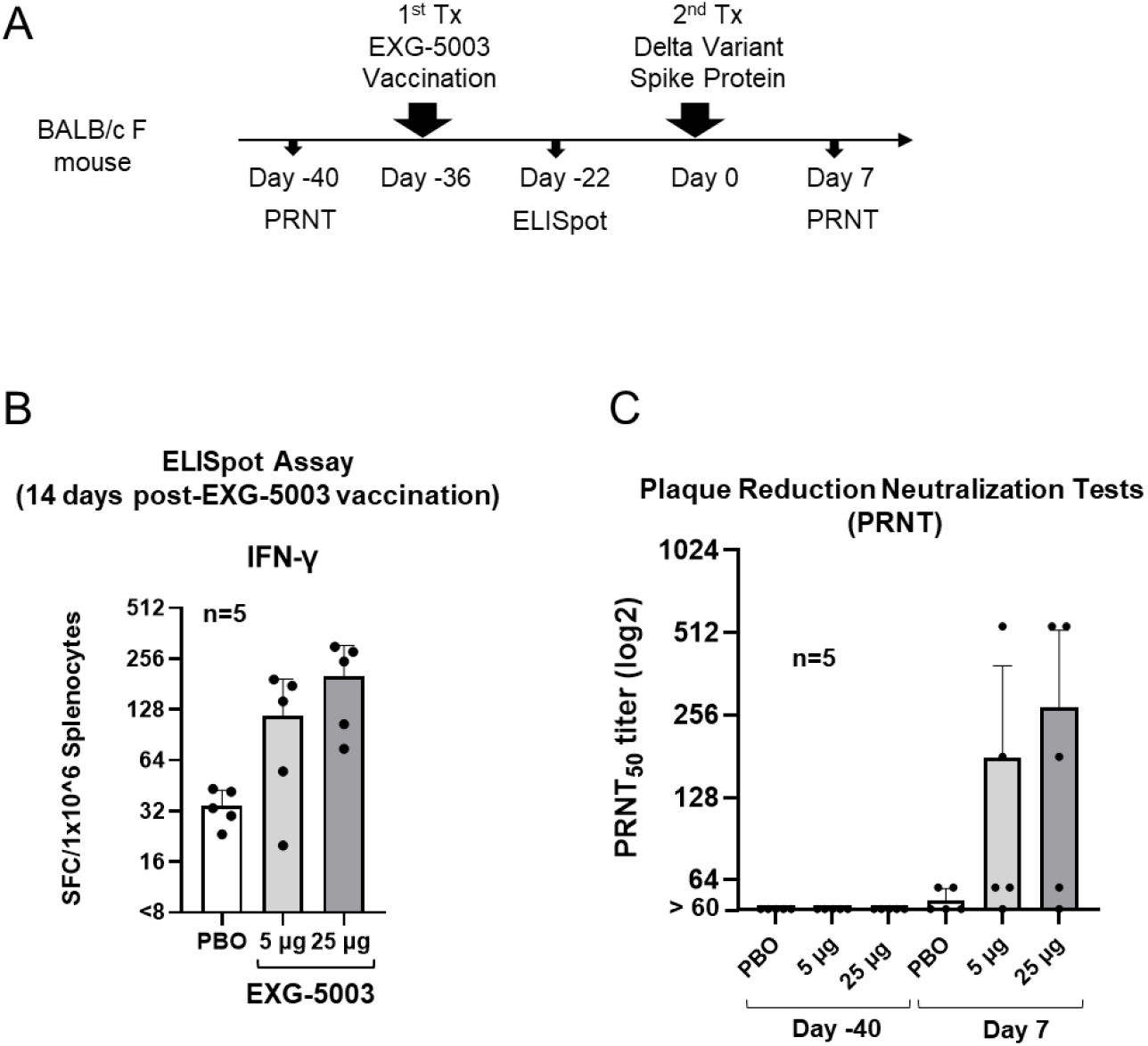
c-srRNA vaccines can prime the immune system for subsequent induction of neutralizing antibodies against a variant antigen upon exposure to the variant antigen. (**A**) A schematic diagram of experimental procedures. On day −40, blood was drawn from female BALB/c mice for a plaque reduction neutralization test (PRNT). On day −36, these mice received an intradermal injection of EXG-5003 RNA, which was formulated as a naked RNA, without a LNP or transfection reagent. On day −22 (14 days after EXG-5003 vaccination), half of mice were sacrificed to obtain splenocytes for ELISpot assays. On day 0, the remaining mice were intradermally injected with the spike protein of the SARS-CoV-2 delta variant (B.1.617.2: R&D Systems) mixed with adjuvant – AddaVax (Invivogen). On day 7 (7 days after the spike protein injection), blood was drawn for the PRNT assay. (**B**) The induction of cellular immunity against the RBD protein by a single intradermal administration of EXG-5003 RNA. The ELISpot assay shows the number of IFN-γ spot-forming cells (SFC) in 1×10^6 splenocytes from immunized mice restimulated by culturing in the presence or absence of a pool of 15mer peptides for RBD of SARS-CoV-2 (original strain). The number of SFC obtained in the presence of peptides was plotted on the graph after subtracting the number of SFC obtained in the absence of peptides (background). The average and standard deviation (error bars) of five mice (n=5) are shown for each group. (**C**) The titer of serum antibodies that can neutralize (50%) the SARS-CoV-2 virus (delta variant B.1.617.2), measured by the PRNT assay. Exposure to the spike protein of the SARS-CoV-2 virus (delta variant B.1.617.2) induced neutralization antibodies against the delta variant of the SARS-CoV-2 virus only in mice vaccinated with EXG-5003, which encodes the RBD of SARS-CoV-2 (original strain).

A plaque reduction neutralization test (PRNT) showed that subsequent exposure of the vaccinated mice to spike protein of SARS-CoV-2 (Delta variant B.1.617.2) induced neutralizing antibodies against the delta variant of SARS-CoV-2 virus in mice vaccinated with EXG-5003 (FIG. 4C). The induction of NAb occurred as early as day 7 after exposure to antigen protein. The rapid induction of neutralizing antibodies indicates that this is a secondary immune response.

Because a c-srRNA vaccine can prime humoral immunity to produce NAb against an antigen varied from the antigen encoded on the c-srRNA vaccine, c-srRNA vaccines can possibly be used as universal vaccines for a range of variants of the pathogen.

### Improvement of the c-srRNA vector

Our initial RNA vector, named c-srRNA1, was based on the TRD strain but contained the TC-83 (A551D) mutation and a shorter 3’-UTR. We reasoned that the exact TRD sequence may work better, because TRD is more resistant to suppression by type I interferons than other VEEV strains, including TC-83 (Spotts et al., 1998). Therefore, we made c-srRNA3, which contains the exact TRD sequence except for the 15-nucleotide addition that confers temperature-controllability. As a wild type control, we also made an srRNA0, which is the same c-srRNA3 vector without the temperature-controllable 15 nucleotides.

We compared the immunogenicity of the three srRNAs using the same RBD protein as an antigen. C57BL/6 mice were treated with intradermal injection of srRNA0-RBD, c-srRNA1-RBD (EXG-5003), or c-srRNA3-RBD, all of which encode the RBD of SARS-CoV-2 [original strain], as naked RNA. As expected, cellular immunity assessed by the presence of antigen-specific IFN-γ-secreting T cells was already induced by day 14 post-vaccination (FIG. 5). The results showed that, as expected, the T-cell response of the c-srRNA1-RBD was stronger than that of the standard, non-temperature-sensitive, non-controllable srRNA-RBD (srRNA0). Furthermore, the results also showed that the T-cell response of the improved version of c-srRNA, i.e., c-srRNA3, was even stronger than that of c-srRNA1 - about a 3-fold improvement over c-srRNA1 and 6-fold improvement compared to the standard srRNA (srRNA0).

**FIG. 5.**
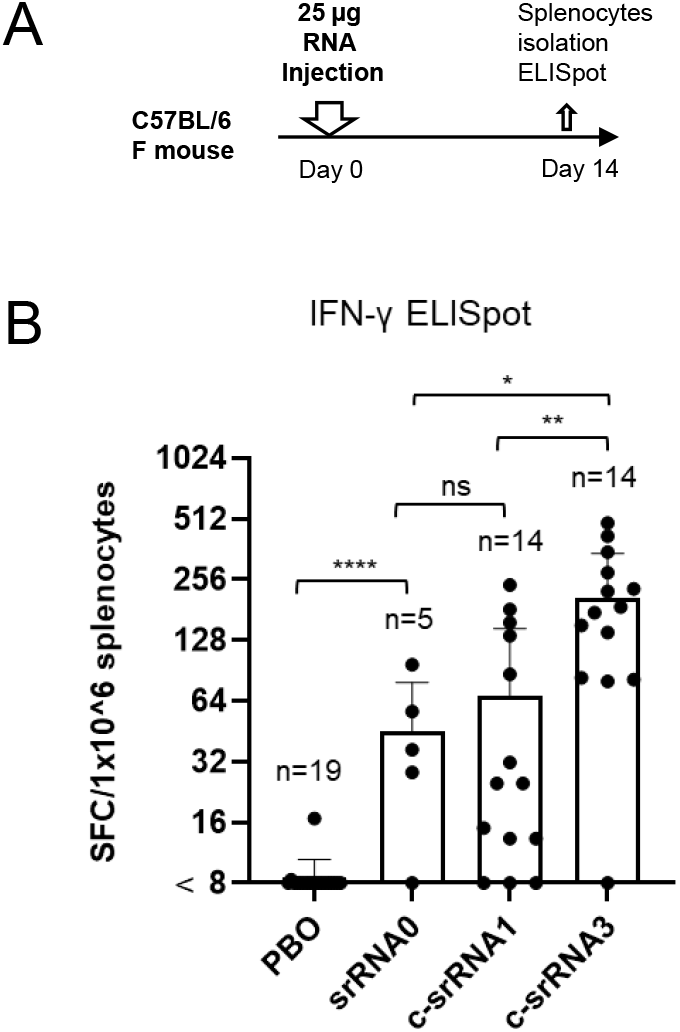
Comparison of srRNA and c-srRNAs for T-cell-inducibility. (**A**) A schematic diagram of experimental procedures. On day 0, mice were intradermally injected with either placebo (PBO, buffer only), srRNA0, c-srRNA1, or c-srRNA3. The srRNA0, c-srRNA1, and c-srRNA3 encode the same RBD of SARS-CoV-2 (original strain). On day 14, mice were sacrificed and splenocytes were isolated for ELISpot assays. (**B**) The ELISpot assay shows the number of IFN-γ spot-forming cells (SFC) in 1×10^6 splenocytes from immunized mice restimulated by culturing in the presence or absence of a pool of 15mer peptides for RBD of SARS-CoV-2 (original strain). The number of SFC obtained in the presence of peptides was plotted on the graph after subtracting the number of SFC obtained in the absence of peptides (background). The average and standard deviation (error bars) are shown for each group.

### EXG-5003 and EXG-5003o boost humoral immunity as a booster vaccine

To test whether a c-srRNA vaccine could be used as a booster vaccine, mice were first vaccinated intradermally with protein (in this case, RBD of SARS-CoV-2 [original strain] and adjuvant). Fourteen days later (Day 14), the mice were further treated with intradermal injection of a placebo (PBO: buffer only); EXG-5003; EXG-5003o (c-srRNA3 encoding the RBD of the omicron variant of SARS-CoV-2 [B.1.1.529 BA.1]); or the RBD protein plus adjuvant (FIG. 6A).

**FIG. 6.**
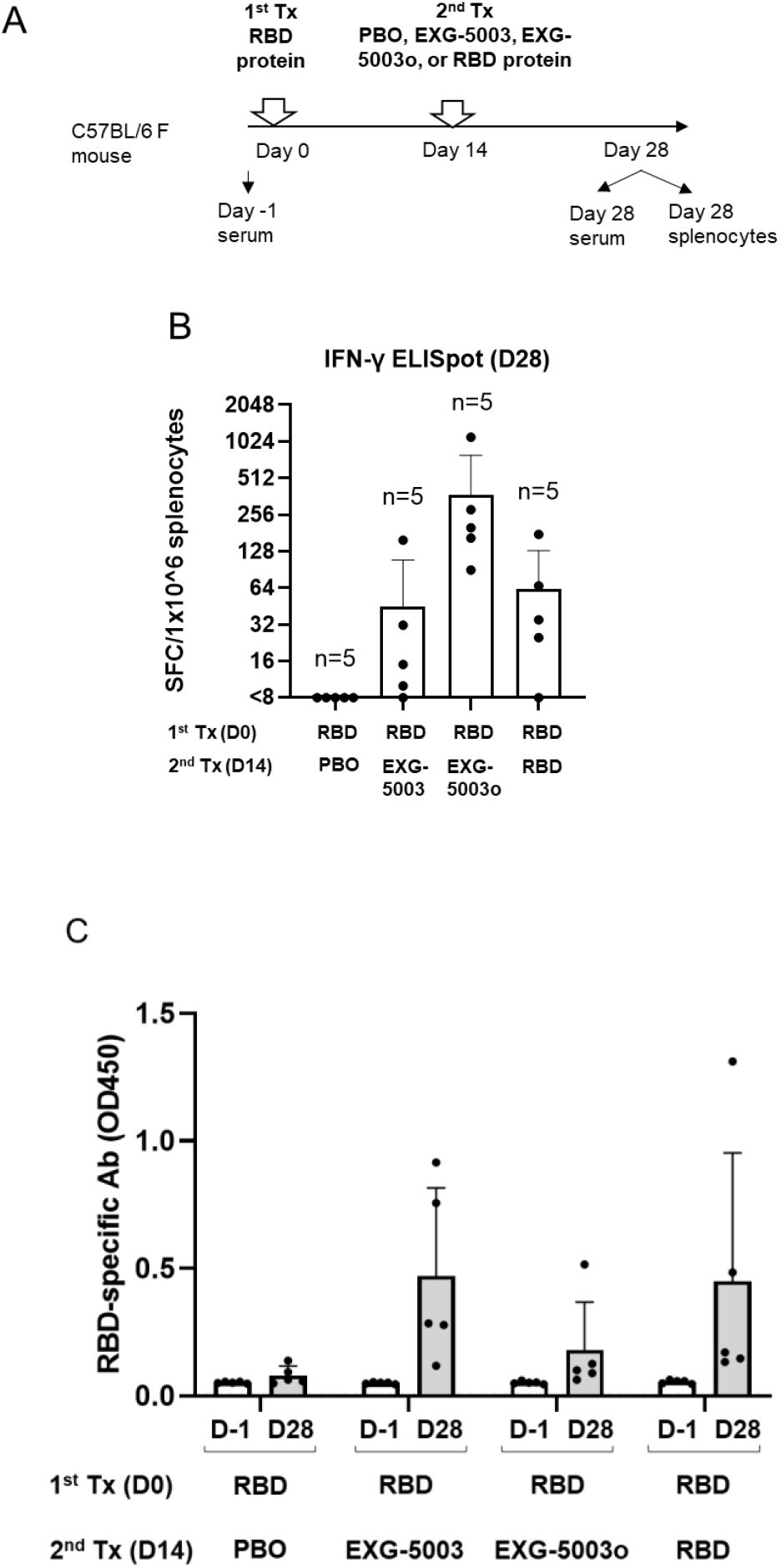
c-srRNA vaccines can be used as a booster to induce antibodies against an antigen, following earlier administration of the antigen protein. **(A)** A schematic diagram of experimental procedures. On day 0 (1^st^ treatment), female C57BL/6 mice were treated with an intradermal injection of 10 μg RBD protein (Sino Biological SARS-CoV-2 [original strain]) + Adjuvant (AddaVax). On day 14 (2^nd^ treatment), the mice were treated with an intradermal injection of a placebo (PBO: buffer only), 25 μg EXG-5003 (c-srRNA1 encoding RBD [original strain]), 25 μg of EXG-5003o (c-srRNA3 encoding RBD [omicron variant B.1.1.529 BA.1]), or 10 μg RBD protein (Sino Biological SARS-CoV-2) + Adjuvant (AddaVax). On day 28, the mice were sacrificed, and splenocytes and serum were collected for ELISpot and ELISA assays. **(B)** The induction of cellular immunity is shown by the frequency of IFN-γ spot-forming cells (SFC) in 1×10^6 splenocytes restimulated by culturing in the presence or absence of a pool of 15mer peptides for RBD of SARS-CoV-2 (original strain) (RBD-PBO, RBD-EXG-5003, RBD-RBD) or 15mer peptides for RBD of SARS-CoV-2 (omicron variant) (RBD-EXG-5003o). The frequency obtained in the presence of peptides is plotted in the graph after subtracting the frequency obtained in the absence of peptides (background). (**C**) The levels of serum antibodies against the RBD protein of the SARS-CoV-2 (original strain), measured by an ELISA assay. The levels of antibodies are represented by the OD450 measurement. The average and standard deviation (error bars) of five mice (n=5) are shown for each group. The data of Day −1 (before the 1^st^ treatment) and the data of Day 28 (after the 2^nd^ treatment) are shown for each group.

On Day 28, cellular immunity was assessed by ELISpot assay using a pool of 15mer peptides for the RBD of SARS-CoV-2 (original strain) for EXG-5003 or 15mer peptides for RBD of SARS-CoV-2 (omicron variant) for EXG-5003o. The RBD (1st) + PBO (2nd) group could not induce cellular immunity, whereas the RBD (1st) + RBD (2nd) group did (FIG. 6B). This suggests that a single dose of the intradermal protein vaccine induces cellular immunity little if at all, but in contrast, a single intradermal injection of c-srRNA vaccine is sufficient to induce cellular immunity. As expected, the RBD (1st) + EXG-5003 (2^nd^) and EXG-5003o (2^nd^) group induced cellular immunity (FIG. 6B).

On Day 28, the levels of serum antibodies against the RBD of the SARS-CoV-2 virus (original strain) were assessed by an ELISA assay (FIG. 6C). The first vaccination with the RBD protein and adjuvant could only weakly induce antibodies. On the other hand, c-srRNA vaccines were able to induce antibodies at a level similar to that achieved by a second vaccination with the adjuvanted protein.

Thus, c-srRNA vaccines can work as booster vaccines for both cellular immunity and humoral immunity.

### c-srRNA vaccine encoding nucleoproteins of SARS-CoV-2 with (EXG-5005) or without (EXG-5004) signal peptide

Our cellular immunity-focused approach allowed us to consider all the proteins (including spike protein) encoded on coronavirus genomes as antigen candidates. Looking for an antigen that would provide broader protection against SARS-CoV, SARS-CoV-2, and MERS-CoV and their variants, we reasoned that the nucleoprotein is most suitable, because (1) nucleoprotein is the most abundant viral protein, followed by the membrane and spike proteins (Finkel et al., 2021); (2) nucleoprotein is overall the most conserved protein among the above indicated Betacoronaviruses (Grifoni et al., 2020); and (3) epitopes for B and T cells are most abundant in the spike and nucleoprotein (Grifoni et al., 2020). This is in line with the earlier proposal that nucleoprotein is the best antigen for a vaccine (Dutta et al., 2020). Notably, a recent report clearly demonstrated that a vaccine using nucleoprotein alone as an antigen can provide spike-independent protective immunity in both hamster and mouse (Matchett et al., 2021).

To test whether nucleoprotein encoded on a c-srRNA vector can itself induce cellular immunity, we generated EXG-5004, a c-srRNA3 encoding a full-length nucleoprotein of SARS-CoV-2 (original strain), and EXG-5005, a c-srRNA3 encoding a fusion protein of CD5 signal peptide and a full-length nucleoprotein of SARS-CoV-2 (original strain) (FIG. 7A). [The two versions, EXG-5004 and EXG-5005, were generated to test whether CD5 signal peptide is required to induce cellular immunity.]

**FIG. 7.**
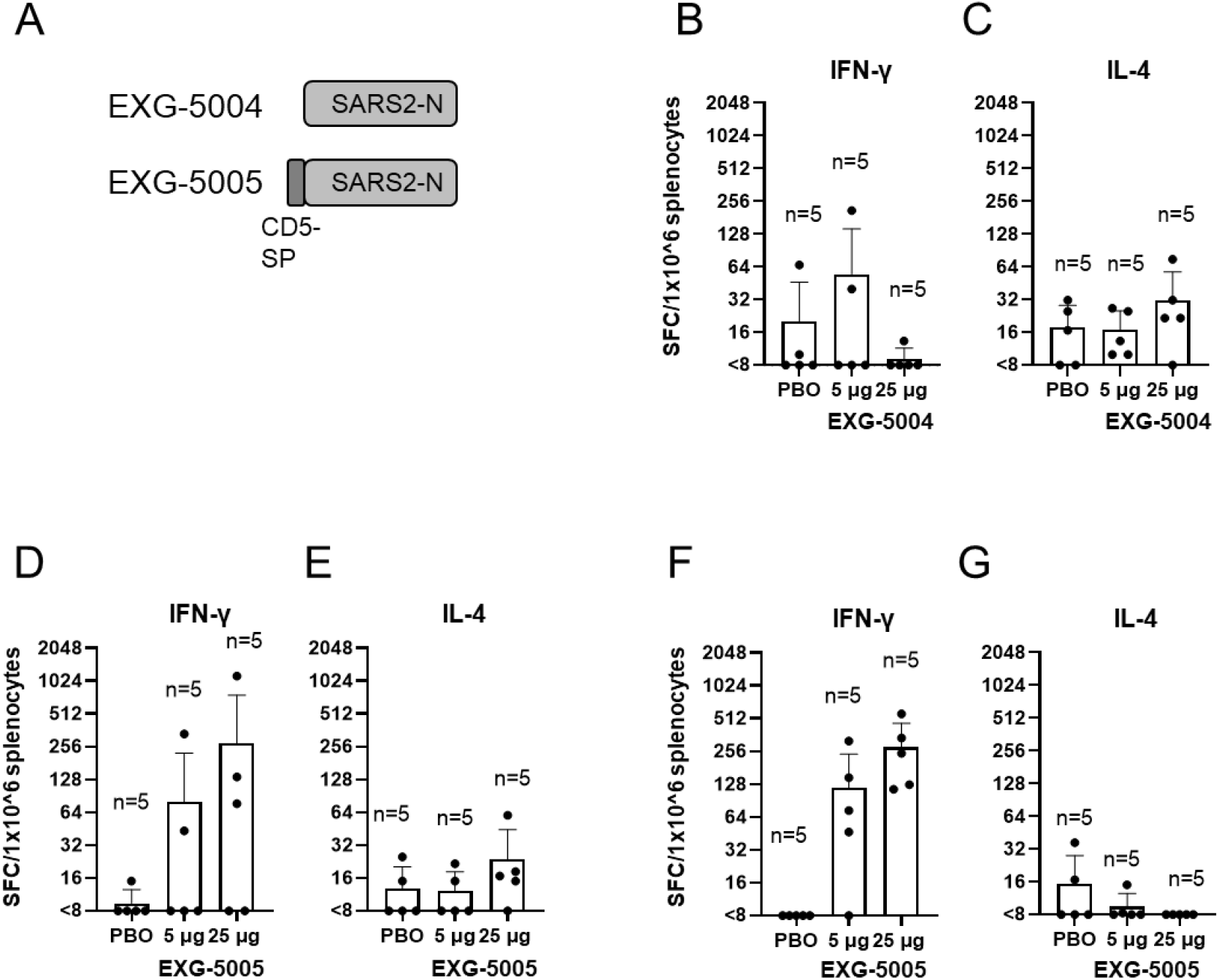
Use of nucleoprotein as an antigen and test for the requirement of signal peptides. (**A**) A schematic drawing of antigen structure. EXG-5004: c-srRNA3 encodes the nucleoprotein (N) protein of SARS-CoV-2 (original strain). EXG-5005: c-srRNA3 encodes a fusion protein of signal peptide sequence of the human CD5 protein and the nucleoprotein (N) protein of SARS-CoV-2 (original strain). (**B - G**) Cellular immunity by ELISpot assays, showing the frequency of IFN-γ spot-forming cells (SFC) or IL-4 SFC in 1×10^6 splenocytes obtained from CD-1 outbred mice or BALB/c inbred mice that were immunized by a single intradermal injection of 100 μL solution containing either 5 μg or 25 μg of EXG-5004, EXG-5005, or a placebo (PBO: buffer only). The splenocytes were restimulated by culturing in the presence and absence of a pool of 15mer peptides for nucleoprotein of SARS-CoV-2 (original strain). The frequency obtained in the presence of peptides is plotted in the graph after subtracting the frequency obtained in the absence of peptides (background). The average and standard deviation (error bars) of five mice (n=5) are shown for each group. (**B**) CD-1 mouse, EXG-5004, IFN-γ. (**C**) CD-1 mouse, EXG-5004, IL-4. (**D**) CD-1 mouse, EXG-5005, IFN-γ. (**E**) CD-1 mouse, EXG-5005, IL-4. (**F**) BALB/c mouse, EXG-5005, IFN-γ. (**G**) BALB/c mouse, EXG-5005, IL-4.

We assessed cellular immunity by ELISpot assays 14 days after vaccinating CD-1 outbred mice by a single intradermal injection of either 5 μg or 25 μg of EXG-5004 or a placebo (PBO: buffer only). As shown in FIG. 7B and 7C, EXG-5004, a nucleoprotein without CD5 signal peptide, induced only weak cellular immunity for both IFN-γ-secreting T cells and IL-4-secreting T cells. Also, there was no response to increasing dose of vaccine (5 μg vs. 25 μg). In contrast, when the CD5 signal peptide was provided, strong induction of IFN-γ-secreting T cells was observed in an antigen-specific manner and in a dose-dependent manner (5 μg vs. 25 μg) (FIG. 7D). As expected, there was only weak induction of IL-4-secreting T cells (FIG. 7D). We infer that the intradermally administered c-srRNA vaccine requires the CD5 signal peptide to induce cellular immunity. The results also indicate that nucleoprotein can be used as an antigen for c-srRNA-induced antigen-specific cellular immunity.

We also tested whether EXG-5005 can induce strong cellular immunity in another mouse strain. Cellular immunity was assessed by ELISpot assays 30 days after vaccinating BALB/c mice with a single intradermal injection of either 5 μg or 25 μg of EXG-5005 or a placebo (PBO: buffer only). As shown in FIG. 7F and 7G, strong induction of IFN-γ-secreting T cells was observed in an antigen-specific and dose-dependent manner. By contrast, there was again weak induction of IL-4-secreting T cells (FIG. 7G). Based on these findings, the induction of cellular immunity and a favorable Th1>Th2 response by intradermal administration of c-srRNA is not mouse-strain specific.

### Generation of EXG-5006: c-srRNA vaccine encoding nucleoproteins of SARS-CoV-2 and MERS-CoV

Our first broad vaccine candidate, EXG-5005, contains nucleoprotein of SARS-CoV-2 (for short, SARS2-N). However, MERS-N shows only 48% identity (Tilocca et al., 2020), and thus, we designed a second vaccine candidate, EXG-5006, that contains a fusion protein of SARS2-N and MERS-N.

Because T-cell epitopes are short linear peptides, we reasoned that nucleoprotein coding sequences from different betacoronavirus strains can simply be fused together and used as a vaccine antigen covering different betacoronaviruses. It is also known that cellular immunity can work for proteins with more variant sequences than B-cell epitopes (which comprise conformational or 3D-structural features). Therefore, we reasoned that a fusion protein of the SARS-CoV-2 nucleoprotein and the MERS-CoV nucleoprotein as a vaccine antigen may provide protection against essentially all betacoronaviruses, including SARS-CoV and variants; SARS-CoV-2 and variants; and MERS-CoV and variants. [SARS-CoV and SARS-CoV-2 nucleoproteins are similar, but the MERS nucleoprotein is very different (Tilocca et al., 2020) (Grifoni et al., 2020)].

To test this notion, we constructed a c-srRNA3 vaccine encoding a fusion protein of the CD5-signal peptide; the SARS-CoV-2 nucleoprotein; and the MERS-CoV nucleoprotein. We generated two versions: non-codon-optimized (natural viral protein sequence) EXG-5006a and the more conventional codon-optimized EXG-5006b (FIG. 8A). We vaccinated BALB/c mice with either EXG-5006a or EXG-5006b by intradermal injection and tested cellular immunity by ELISpot assays. Interestingly, the codon-optimized version was less effective than the non-optimized version (Supplemental FIG. S3). Therefore, we continued to use the natural protein sequence without codon optimization. As expected, intradermal administration of EXG-5006a induced cellular immunity against both the SARS-CoV-2 nucleoprotein (FIG. 8B) and MERS-CoV nucleoprotein (FIG. 8D).

**FIG. 8.**
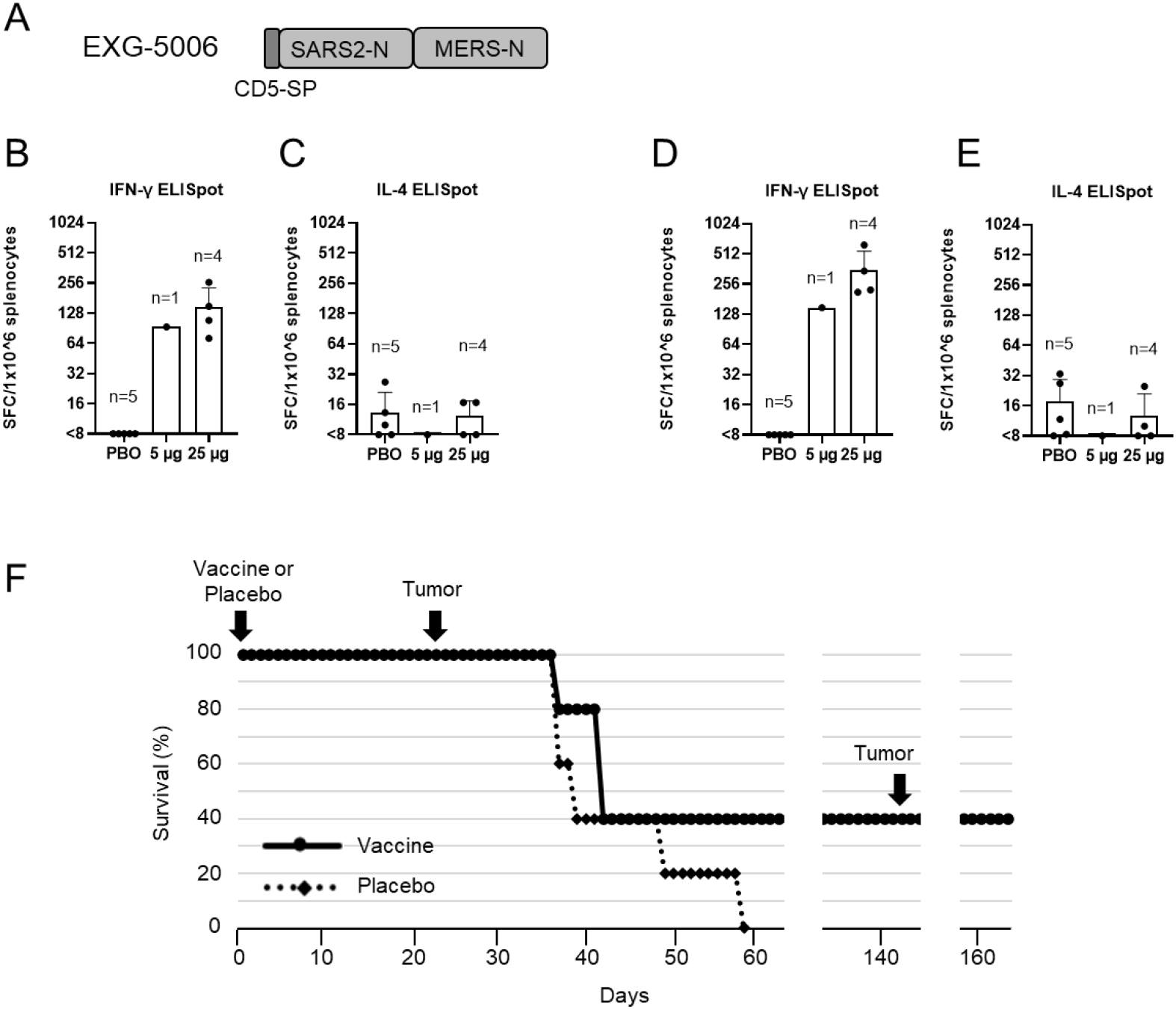
EXG-5006, encoding a fusion protein of SARS-CoV-2 and MERS-CoV nucleoproteins, can eliminate cells expressing those antigens in mouse. (**A**) A schematic drawing of EXG-5006, encoding a fusion protein of CD5 signal peptide, nucleoprotein of SARS-CoV-2, and a nucleoprotein of MERS-CoV. (**B - E**) Cellular immunity by ELISpot assays, showing the frequency of IFN-γ (**B, D**) spot-forming cells or IL-4 (**C, E**) spot-forming cells in 1×10^6 splenocytes obtained from BALB/c mice on Day 14 after vaccinating by a single intradermal injection of 100 μL solution containing either 5 μg (n=1) or 25 μg (n=4) of EXG-5006, or a placebo (PBO: buffer only: n=5). The splenocytes were restimulated by culturing in the presence and absence of a pool of 15mer peptides for nucleoprotein of SARS-CoV-2 (original strain) (**B, C**) or a pool of 15mer peptides for nucleoprotein of MERS-CoV (**D, E**). The frequency obtained in the presence of peptides is plotted in the graph after subtracting the frequency obtained in the absence of peptides (background). (**F**) shows the survival (%) of female BALB/c mice vaccinated with EXG-5006, followed by injection of 4T1 tumor cells expressing the same antigen (A fusion protein of nucleoproteins of SARS-CoV-2 and MERS-CoV, without CD5 signal peptides).

In vaccines targeting a cellular immunity mechanism to infectious agents, the vaccine is expected to eliminate infected cells via the action of antigen-specific CD8+ cytotoxic T cells. To model cells infected with a virus, we used the 4T1 breast cancer cell line derived from a BALB/c mouse. When injected into a BALB/c mouse, 4T1 cells grow rapidly and form tumors. This syngeneic mouse model was used as a proxy for the rapid increase of infected cells. 4T1 cells expressing a fusion protein of the <u>S</u>ARS-CoV-2 nucleoprotein and the <u>M</u>ERS-CoV nucleoprotein (named 4T1-SMN cells) were established by transfecting a plasmid vector encoding the fusion protein under the CMV promoter, so that the fusion protein is constitutively expressed. This fusion protein is the same as the antigen for EXG-5006, but the CD5 signal peptide was removed from the N-terminus of the protein.

BALB/c mice were first vaccinated by intradermal administration of EXG-5006a. Subsequently, 4T1-SMN cells were injected into the BALB/c mice on day 24 (24 days post-vaccination) (FIG. 8D). As expected, 4T1-SMN cells grew rapidly in mice that received a placebo (non-vaccinated group). On the other hand, the growth of 4T1-SMN tumors was suppressed in the mice that received EXG-5006a vaccination. Two mice received 25 μg of EXG-5006a vaccine, and although the tumor grew initially, the mice eventually became tumor-free and survived. Furthermore, even after a second round of injection of 4T1-SMN cells on day 143 after the vaccination, no tumors grew, and the mice were tumor-free and continued to live (FIG. 8D). The results indicate that the EXG-5006a vaccine induces strong cellular immunity and eliminates cells expressing the nucleoproteins of SARS-CoV-2 and MERS-CoV.

### Generation of pan-coronavirus booster vaccine EXG-5008: a c-srRNA vaccine encoding RBDs and nucleoproteins of SARS-CoV-2 and MERS-CoV

Given all the data thus far, it is conceivable to use c-srRNA as a vaccine tailored both to induce cellular immunity and to prime or boost humoral immunity against essentially all betacoronaviruses. In the current SARS-CoV-2 pandemic, most of the global population is either already exposed, or vaccinated with vaccines targeting the RBD or spike proteins. In a booster vaccine scenario, a c-srRNA encoding the RBD, as shown here for EXG-5003 and EXG-5003o (encoding the RBD of omicron variant), would stimulate cellular immunity via T-cell epitopes against the RBD and restore NAb titer against the RBD. In addition, including the more evolutionarily conserved nucleoprotein would extend cellular immunity via CD8+ cytotoxic T cells. Therefore, we designed and generated a c-srRNA booster vaccine (called EXG-5008) encoding a fusion protein comprising the CD5 signal peptide, RBD of SARS-CoV-2, nucleoprotein of SARS-CoV-2, nucleoprotein of MERS-CoV, and RBD of MERS-CoV (FIG. 9A).

**FIG. 9.**
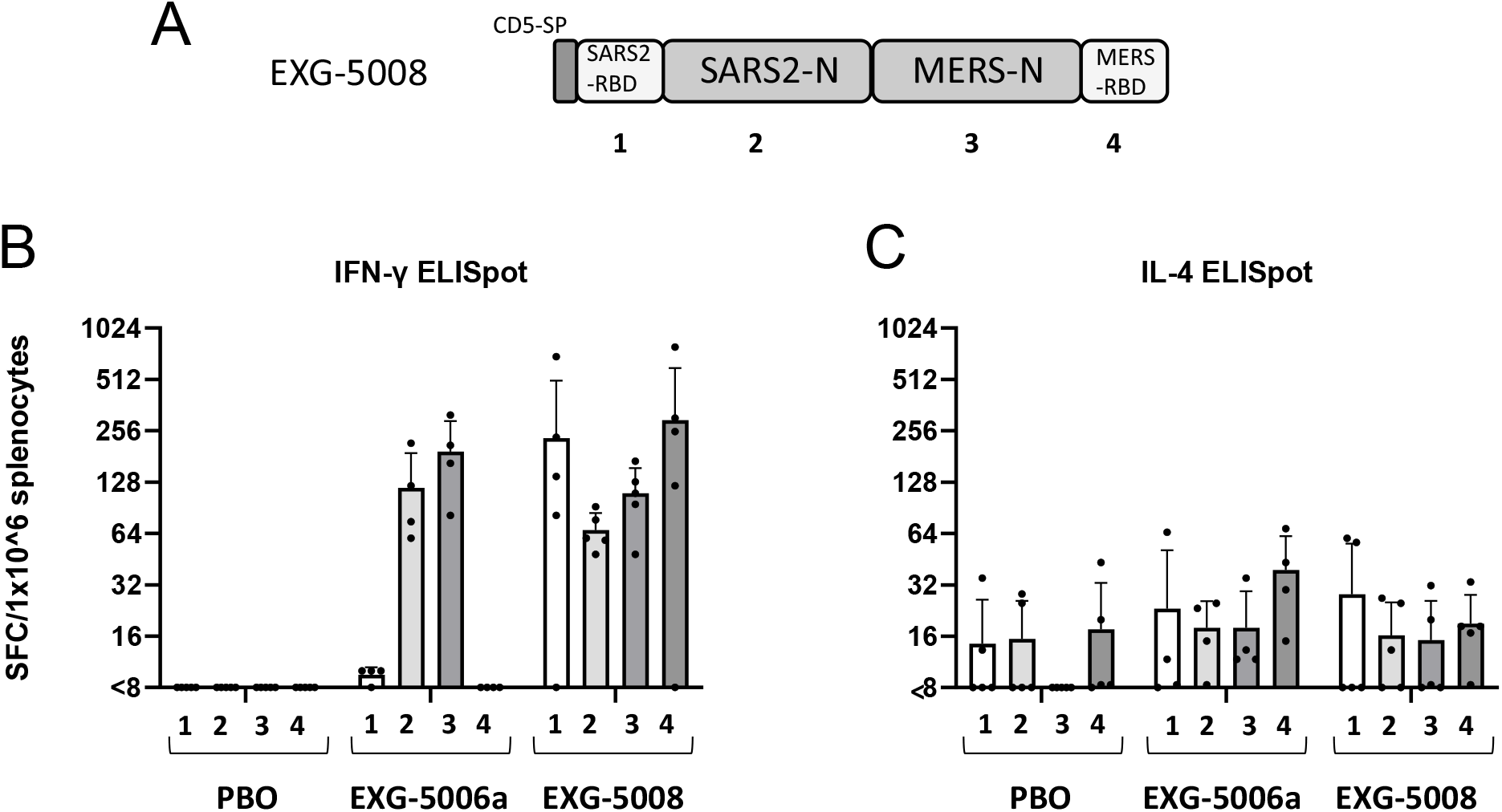
Generation of EXG-5008, encoding RBD and nucleoproteins of SARS-CoV-2 and MERS. (**A**) A schematic diagram of EXG-5008, encoding a fusion protein of the signal peptide of CD5, the RBD and nucleoprotein of SARS-CoV-2, and the RBD and nucleoprotein of MERS-CoV. (**B, C**) The frequency of IFN-γ spot-forming cells (**B**) and the frequency of IL-4 spot-forming cells (**C**) in 1×10^6 splenocytes obtained from female C57BL/6 mice that were immunized by a single intradermal injection of 100 μL solution containing either placebo (PBO: buffer only), 25 μg of EXG-5006a, or 25 μg of EXG-5008. The splenocytes were restimulated by culturing in the presence or absence of pools of peptides: 15mer peptides for RBD of SARS-CoV-2 (original strain) (**1**); 15mer peptides for nucleoprotein of SARS-CoV-2 (original strain) (**2**); 15mer peptides for nucleoprotein of MERS-CoV (**3**); 15mer peptides for spike of MERS-CoV (**4**). The cellular immunity was analyzed by ELISpot assays. The frequency obtained in the presence of peptides is plotted in the graph after subtracting the frequency obtained in the absence of peptides (background). The average and standard deviation (error bars) of five mice (n=5) for PBO, four mice (n=4) for EXG-5006, and five mice (n=5) for EXG-5008, are shown for each group. Splenocytes were isolated 14 days after vaccination.

To test this vaccine, mice were vaccinated with intradermal injection of a placebo (PBO: buffer only); EXG-5006a; or EXG-5008. On day 14 post-vaccination, cellular immunity was assessed by ELISpot assays. As shown in FIG. 9B, EXG-5008 stimulated cellular immunity against all the proteins encoded in this vaccine construct: RBD of SARS-CoV-2, nucleoprotein of SARS-CoV-2, nucleoprotein of MERS-CoV, and RBD of MERS-CoV. As expected, EXG-5006a stimulated cellular immunity only against the nucleoprotein of SARS-CoV-2 and the nucleoprotein of MERS-CoV.

## Discussion

Currently available SARS-CoV-2 vaccines focus on inducing NAb against the spike protein or RBD. However, this approach encounters two challenges: small amino acid changes among variants often cause conformational changes of the protein that could significantly reduce NAb effectiveness, and subunit vaccines against pathogens generally do not provide long-lasting pathogen-specific antibody production, so that frequent booster vaccination is required. Ideally, future vaccines should broadly address a range of variants of SARS-CoV-2, but also other betacoronaviruses such as SARS-CoV and MERS-CoV (Dolgin, 2022).

Broad specificity is a very tall order for vaccines based on the spike protein of SARS-CoV-2 as an antigen, because the spike protein is not well conserved among betacoronaviruses - even between SARS-CoV and SARS-CoV-2 (as detailed below). In this report, we have shown that these challenges can potentially be met by the combination of intradermal administration of a c-srRNA vaccine backbone and a unique antigen design – a fusion protein of viral surface proteins (RBD) and non-surface proteins (nucleoprotein) from evolutionarily diverged SARS-CoV-2 and MERS-CoV (FIG. 9 and FIG. 10).

**FIG. 10.**
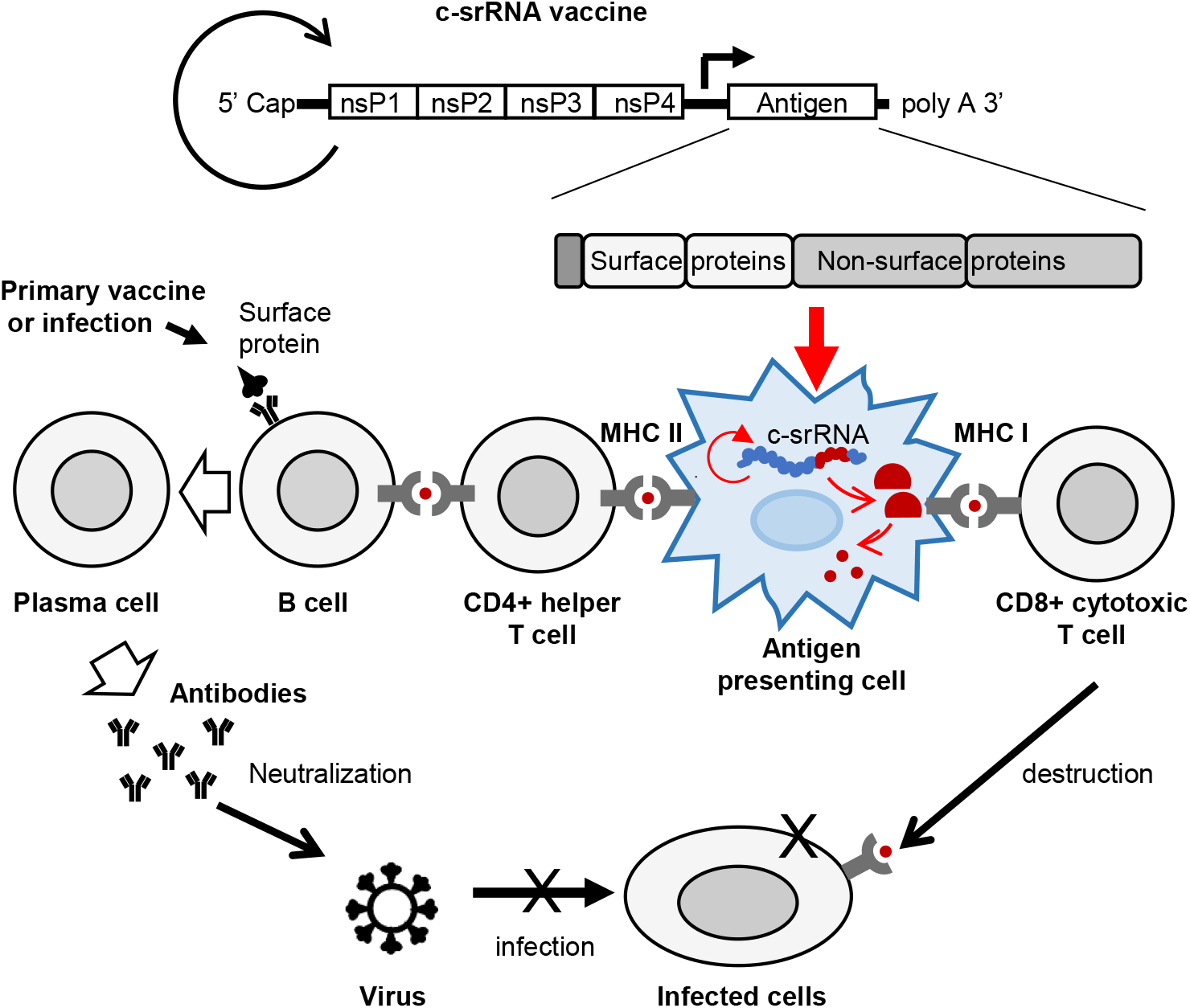
A model showing how c-srRNA booster vaccine works. Primary series of vaccines or prior infections expose naïve B cells to viral surface proteins and turn them into memory B cells in a manner dependent on CD4+ helper T cells. Skin delivery of a c-srRNA booster vaccine encoding a fusion antigen of the viral surface proteins and non-surface proteins, whose sequences are more evolutionarily conserved than the surface proteins, is primarily incorporated into skin antigen presenting cells such as Langerhans cells and dendritic cells. Within the antigen presenting cells, c-srRNA replicates and produces the fusion antigen. The antigen presenting cells digest the antigen into peptides, and present these peptides to T cells. The peptides presented through this pathway stimulate MHC-I-restricted CD8+ cytotoxic T cells as well MHC-II-restricted CD4+ helper T cells. CD8+ cytotoxic T cells eliminate infected cells. CD4+ helper T cells stimulate the memory B cells and enhance or restore the production of neutralizing antibody, which prevents infection.

The c-srRNA platform incorporates the benefits of mRNA utilization, including the absence of genome integration, rapid development and deployment, and a simple manufacturing process, as well as the additional advantages of srRNA platforms such as longer expression (Johanning et al., 1995) and no use of modified nucleotides (reviewed in (Pushko and Tretyakova, 2014)(Brito et al., 2015)(Lundstrom, 2016)(Blakney et al., 2021)). c-srRNA then adds the following features:

1. suitability for intradermal administration, with its optimal expression at skin temperature of about 30-35°C, where antigen expression and T cell immunogenicity is about 6-fold stronger than the commonly used srRNA injected intradermally;
2. potentially greater safety due to inactivation at core body temperature of about 37 °C, reducing the risk caused by any systemic distribution and uncontrollable RNA replication and providing a potential “safety switch” by skin warming at the site of administration if any local adverse events occur;
3. formulation as naked RNA in Lactated Ringer’s solution, without lipid nanoparticles or any other transfection reagents that provide no or only a several-fold increase of gene of interest expression or immunogenicity in intradermal administration (Golombek et al., 2018)(Johansson et al., 2012)(Blakney et al., 2019) and run the risk of skin irritation and allergic reactions to nanoparticles; in fact, this is the only COVID-19 RNA vaccine that does not use nanoparticles (Kisby et al., 2021);
4. capacity to induce cellular immunity via MHC-I-restricted CD8+ cytotoxic T cells as well as MHC-II-restricted CD4+ helper T cells in an antigen-specific manner, most likely resulting from its uptake by skin APCs (Blakney et al., 2019), in which it replicates and produces antigen. The antigen is digested into peptides (T cell epitopes) and presented to T cells in local skin regions or lymph nodes;
5. lack of appreciable induction of humoral immunity, i.e., antibody production, thus achieving a “pure” T-cell-inducing vaccine; this may be due to factors including the lack of B cell stimulation from the paucity of B cells in the skin (Blakney et al., 2019); the lack of circulation of antigen protein into lymph nodes to encounter B cells; and/or the lack of proper presentation of the three-dimensional structure of the antigen; and
6. capacity to induce humoral immunity, i.e., enhancing antibody production, either when the antigen protein is administered after c-srRNA or when the antigen protein is administered before c-srRNA.

In this report, we have explored different antigen designs using betacoronaviruses as a model. The unique features of the c-srRNA vaccine platform allow for an optimized design of a potential pan-coronavirus vaccine. Considering that many people have already been exposed to the spike protein or RBD of SARS-CoV-2 via infection or vaccinations, EXG-5008, which encodes a fusion protein of a CD5 signal peptide, RBD and nucleoprotein of SARS-CoV-2, and RBD and nucleoprotein of MERS-CoV, can be considered as a pan-coronavirus booster vaccine candidate. EXG-5008’s specific combination of antigens and its intradermal application provide the following features:

1. The antigens encoded on the c-srRNA construct do not directly stimulate B cells, and thus this vaccine alone does not induce humoral immunity. Although this is a disadvantage for a vaccine used in a primary vaccination series against an infectious agent, it is an advantage for multivalent antigen designs, because, unlike traditional vaccines, it does not depend in part on features of the three-dimensional structure of antigens, making it feasible to use a fusion protein combining different antigens.
2. Notwithstanding the foregoing feature, when c-srRNA encoding the RBD was used as a booster vaccine following a vaccine (recombinant RBD with adjuvant) that can prime or induce humoral immunity, it was able to stimulate cellular immunity, which increased the level of antibodies or NAb against RBD. The spike proteins of SARS-CoV-2 and SARS-CoV share 76% of their sequence, including many T cell epitopes (Grifoni et al., 2020). Therefore, EXG-5008 should be effective as a booster for SARS-CoV-2, SARS-CoV, and their variants. On the other hand, the spike proteins of SARS-CoV-2 and MERS-CoV are much less alike, sharing only 35% sequence similarity (Grifoni et al., 2020); and therefore, EXG-5008 also encodes the RBD of MERS-CoV. Overall, EXG-5008 is designed to be effective as a booster for SARS-CoV-2, SARS-CoV, MERS-CoV, and their variants – in other words, a wide range of betacoronaviruses. It is also relevant that many T cell epitopes, often specific to certain MHC haplotypes, are present in each single protein, further reducing the probability of one or even several mutations causing an extensive loss of immunogenicity. By contrast, traditional vaccines that induce a B cell response mainly rely on the three-dimensional structure of the protein, and thus even a single mutation may substantially alter conformation of the protein, which may lead to a loss of immunogenicity.
3. Furthermore, EXG-5008 also encodes the nucleoproteins of SARS-CoV-2 and MERS-CoV, providing an additional route to strong cellular immunity against both. The nucleoproteins of SARS-CoV-2 and SARS-CoV share 90% of their sequence (Grifoni et al., 2020), so that once again, EXG-5008 is designed to provide strong cellular immunity against SARS-CoV-2, SARS-CoV, and their variants. Like the spike proteins, the nucleoproteins of SARS-CoV-2 and MERS-CoV are far less similar, with only 48% sequence similarity (Grifoni et al., 2020); and therefore, EXG-5008 also encodes the nucleoprotein of MERS-CoV. Reinforcing its action against the spike protein, the combination of nucleoprotein antigens in EXG-5008 not only increases the stimulation of cellular immunity but also further decreases the likelihood that any particular mutation will promote escape of the pathogen from vaccine protection.
4. c-srRNA’s reliance on short peptide epitopes for cellular and humoral immunogenicity also provides advantages for more general c-srRNA application toward universal vaccines. The concept of a fusion antigen of viral surface proteins for NAb production and evolutionarily conserved non-surface proteins for cellular immunity may be widely applicable to many infectious pathogens (FIG. 10).

In summary, the results thus far are consistent with the concept of a “dendritic cell” vaccine specifically targeting antigen presenting cells. Cellular immunity is known to provide long-lasting protection from severe illness, hospitalization, and death. And when used as a booster vaccine, we expect that EXG-5008, with its unique combination of the c-srRNA platform and fusion antigen design, would boost NAb levels and also provide cellular immunity against a wide range of betacoronaviruses. In addition, although the lack of primary stimulation of humoral immunity is a drawback for initial vaccine administration against infectious diseases, it can be a favorable targeting feature of c-srRNA for other potential applications – particularly for possible therapeutic cancer vaccines, which specifically and only require cellular immunity.

## Acknowledgments

We thank David Schlessinger, Dan L. Longo, Thomas White, Stephen Feinstone for discussion and critical reading of the manuscript. We thank Satomi Amano and Seraina Aguilar for technical help. We thank the members of Elixirgen Therapeutics for discussion, support, and encouragement. This work was supported by Elixirgen Therapeutics. FACS-ICS work was supported in part by the Intramural Program of the National Institute on Aging, National Institutes of Health.

## Author contributions

MSHK conceived and supervised the project, designed c-srRNA vectors and antigens, designed experiments, and analyzed the data. TA, MA, and MB performed mouse studies. HY and PR performed molecular biology work. EL performed ELISpot assays. JK and HY performed ELISA assays. JK and ACK performed cell culture work. NPW and AA performed FACS-ICS and analyzed the data. EK and YL provided MicronJet600 device and training for its usage. MSHK wrote the manuscript with help from ACK, TA, HY, and NPW.

## Materials and Methods

### Vector construction

Most self-replicating RNA (srRNA), also called self-amplifying RNA (saRNA or SAM), are based on the Venezuelan Equine Encephalitis virus (VEEV), where a subgenomic region encoding structural proteins is replaced with a gene of interest (reviewed in (Pushko and Tretyakova, 2014)(Brito et al., 2015)(Lundstrom, 2016)(Blakney et al., 2021)). Two VEEV strains have been widely used: Trinidad donkey (TRD) strain and its attenuated strain TC-83 that accumulated 12 genomic mutations (Kinney et al., 1989). A commonly used vector backbone is a VEEV-TRD with slight sequence modifications (A551D and P1308S) (Petrakova et al., 2005)(Yoshioka et al., 2013)(Mckay et al., 2020). More specifically, we started with a plasmid T7-VEE-GFP, assembled using synthesized DNA fragments based on publicly available sequence information (Yoshioka et al., 2013).

To produce an srRNA that optimally functions around 33°C (skin temperature), we systematically mutated the non-structural proteins of our initial RNA vector and tested the expression levels of the gene of interest encoded in the sub-genomic region at 33°C and 37°C. As a guide, we used a published database of a total of 7480 mutants that were produced by VEEV viral replication at 30°C or at 40°C (Data Set S1 from (Beitzel et al., 2010)). We identified a mutant that functions at 30-35°C but is inactivated at ≥37°C. The mutant has a five amino acid (TGAAA) insertion between 586 (N)-587 (T) of the nsP2 protein. This vector was called T7-VEE-GFP-ts. Subsequently, one “G” nucleotide immediately after the T7 promoter sequence was deleted, so that the *in vitro* transcribed mRNA has an authentic 5’-end of the alphavirus and can be 5’-capped by TriLink BioTechnologies’ CleanCap Technology (Henderson et al., 2021). Furthermore, to remove the unnecessary IRES-PAC, an AflII-SphI DNA fragment was deleted, resulting in the truncated shorter 3’-UTR. This RNA vector was called c-srRNA1-GFP.

As shown in Supplemental Data, c-srRNA1-GFP showed stronger GFP expression than T7-VEE-GFP-ts2 and a synthetic modified mRNA encoding GFP (with 5-methylcytidine and pseudouridine modifications) (Warren et al., 2010)). Also, both srRNAs showed GFP expression for a longer time than a synthetic modified mRNA. The synthetic modified mRNA was expressed in both 33°C and 37°C conditions, whereas both T7-VEE-GFP-ts2 and c-srRNA1-GFP were not expressed at 37°C, as designed.

### Further Improvement of the c-srRNA vector

Our initial c-srRNA1 was based on the TRD strain but contained the VEEV TC-83 mutation (A551D) and also a shorter 3’-UTR. We reasoned that the exact TRD sequence may work better, because TRD is more resistant to suppression by type I interferons than other VEEV strains such as TC-83 (Spotts et al., 1998). Therefore, we made c-srRNA3, which is the exact same sequence as the TRD sequence, except for the 15-nucleotide addition for temperature-controllability. As a wildtype control, we also made an srRNA0, which is exactly the same c-srRNA3 vector, but without the temperature-controllable 15 nt.

### Selection of RBD as an antigen for EXG-5003

When we selected the RBD as the antigen in early 2020, there was no published report on a vaccine against SARS-CoV-2. Therefore, the RBD was selected as the antigen for the following reasons: For the study of SARS-CoV (or SARS-CoV-1), the RBD domain of the spike protein of SARS-CoV had been identified as a key domain for binding to cellular angiotensin converting enzyme 2 (ACE2)-receptor and a suitable target for drugs and vaccines (Du et al., 2009). At the time of our antigen design, it was reported that the amino acid sequence of SARS-CoV-2 RBD is well conserved: among the 1,609 SARS-CoV-2 strains completely sequenced then; only 32 strains contained amino acid mutations in the RBD (Ou et al., 2021). Animal studies had already demonstrated that vaccination with the RBD can provide immune protection against SARS-CoV (Du et al., 2009). While prospective vaccines using the full spike protein had been implicated in liver toxicity (Weingartl et al., 2004), the RBD alone has not been linked to any adverse events in these animal studies (Du et al., 2009). Therefore, vaccination in terms of safety and efficacy, the RBD from SARS-CoV-2, isolated from Wuhan (NCBI accession number: NC_045512, (Wu et al., 2020)), was the logical choice for an antigen at that point.

### Viral Genes/proteins used in this study

- RBD (also called SARS-RBD, original strain): Spike receptor binding domain of Severe acute respiratory syndrome coronavirus 2 isolate Wuhan-Hu-1 (NCBI accession number: NC_045512).
- RBD (omicron variant): Spike receptor binding domain of Severe acute respiratory syndrome coronavirus 2, omicron variant [B.1.1.529 BA.1]. Amino acid sequence of RBD (original strain) was replaced by G339D, S371L, S373P,S375F, N440K, G446S, S477N, T478K, E484A, Q493R, G496S, Q498R, N501Y, Y505H.
- SARS2-N (original strain): nucleoprotein of Severe acute respiratory syndrome coronavirus 2 isolate Wuhan-Hu-1 (NCBI accession number: NC_045512).
- MERS-N: nucleoprotein of Middle East respiratory syndrome-related coronavirus (MERS), strain HCoV-EMC (RefSeq accession number: GCF_000901155.1)
- MERS-RBD: Spike receptor domain of Middle East respiratory syndrome-related coronavirus (MERS), strain HCoV-EMC (RefSeq accession number: GCF_000901155.1).
- CD5sp: Signal peptide sequence from homo sapiens CD5 molecule, transcript variant 1 (NCBI accession number: NM_014207.4).

### Preparation of c-srRNA

All c-srRNAs were produced by *in vitro* transcription. NEB 10-beta Competent E. coli (C3019H/C3019I) was transformed with a plasmid DNA using High Efficiency Transformation Protocol (New England Biolabs). After picking a single colony, E. Coli carrying the plasmid DNA was cultured in Luria Broth (LB) with 100 μg/mL ampicillin. Plasmid DNA was purified using HiSpeed Plasmid Midi Kit (Qiagen). Plasmid DNA was linearized by MluI. *In vitro* transcription (IVT) of c-srRNA with Cap1 and polyA was performed using *in vitro* transcription of a plasmid DNA with Cleancap AU (Trilink) according to the manufacturer’s protocol. Reagents used for the IVT were the following: MluI-linearized DNA Template, Nucleoside-5’-Triphosphate (NTP) Set (TriLink cat. no. N-1505), T7 RNA polymerase (New England BioLabs cat. no. M0251S), Yeast Inorganic Pyrophosphatase (New England BioLabs cat. no. M2403S), Murine RNase Inhibitor (New England BioLabs cat. no. M0314S), 1M Tris-HCL (pH 8.0), RNase Free (Thermo Fisher Scientific cat. no. AM9856), Dithiothreitol (DTT) (Sigma cat. no. 43816), Spermidine (Sigma Aldrich cat. no. 85558-1G), Triton X-100 (VWR cat. no. 80503-490), 1M Magnesium Acetate (Sigma Aldrich cat. no. 63052), UltraPure™ DNase/RNase-Free Distilled Water (ThermoFisher Scientific cat. no. 10977015).

IVT reactions for a total volume of 100 μL were assembled in 200 μL PCR tube in the following order: RNase free water, NTPs, CleanCap AU, 10X Transcription Buffer, DNA template, Murine RNase Inhibitor, Yeast Inorganic Pyrophosphatase, and T7 RNA Polymerase.

The reaction was performed at 37°C for 3 hours, followed by the addition of 5 μL RQ1 RNase-Free DNase (Promega, Cat# M6101) and incubation at 37°C for 15 minutes. Subsequently, RNAs were purified using MEGAclear™ Transcription Clean-Up Kit from ThermoFisher Scientific (Cat#:AM1908) according to the manufacturer’s protocol. In brief, 350 μL of binding buffer and 250 μL of 100% ethanol to each IVT reaction tube, which are now applied to MEGAclear column. After centrifugation at 10,000 rpm for 1 minute and discarding the flow through, the column was washed twice by 500 μL wash buffer. RNA was eluted from the column by two times applications of 60 μL 1 mM sodium citrate, preheated at 95°C, followed by centrifugation at 13,000 rpm for 1 min, and one time application of 60 μL UltraPure DNase/RNase-free distilled water, preheated at 95°C, followed by centrifugation at 13,000 rpm for 1 min. Finally, 3 tubes of eluted RNAs were combined into one tube, resulted in purified RNAs in 180 μL of 0.67 mM sodium citrate. RNA solutions are usually stored at −80°C.

### Preparation of synthetic modified mRNA

Synthetic modified mRNA-GFP (with 5-methylcytidine and pseudouridine modifications) was prepared according to a published protocol (Warren et al., 2010). Synthetic modified mRNA-LUC (with 5MoU modifications) was purchased from TriLink Technologies.

### Experimental Animals

C57BL/6, BALB/c, and CD-1 mice were purchased from Charles River. 5 mice were housed in each individually ventilated cage in a temperature regulated room with a 12:12 light/dark cycle. They were allowed *ad libitum* access to food and water. All mice were acclimatized for at least 72 hours before the start of the experiment. All experiments were approved by the Institutional Animal Care and Use Committee of Elixirgen Therapeutics (Protocol #: ET-IAC-001).

### Cell Culture

Mouse mammary tumor cell line, 4T1 (ATCC, CRL-2539) and 4T1-Luc2 (ATCC, CRL-2539-Luc2), were purchased from ATCC. Human adult dermal fibroblast cells (HDFa) were purchased from ThermoFisher Scientific. HDFa cells were transfected with mRNAs using

Lipofectamine MessengerMAX (ThermoFisher Scientific) and cultured in the DMEM supplement with 500 ng/ml B18R (ThermoFisher Scientific).

### Injection of c-srRNA into mouse skin

Mice were randomly divided into groups and the fur on their hindlimbs was shaved to expose the skin one day before injection. 5 μg or 25 μg of c-srRNA reconstituted in Lactated ringer’s (LR) solution were intradermally (i.d.) injected with MicronJet600 (NanoPass Technologies, Israel) onto the shaved skin.

### Live animal imaging

Bioluminescence imaging was performed using an Ami HT imaging system (Spectral Instruments Imaging) and analysis software (Aura imaging). Briefly, mice were injected with 150 mg/kg D-luciferin (PerkinElmer) intraperitoneally and placed in a chamber for isoflurane anesthesia. Mice were then placed in the Ami HT imaging system and monitored for bioluminescence within 10 min after D-luciferin injection.

### Injection of proteins

The following recombinant proteins were purchased and mixed with adjuvant (AddaVax, Invivogen) before injection into mice.

- RBD protein of SARS-CoV-2 (original strain): Sino Biological, SARS-CoV-2 [2019-nCoV] RBD-His Recombinant Protein (catalog number #40592-V08B).
- Spike protein of SARS-CoV-2 (B.1.617.2, also called delta variant): R&D Systems, Recombinant SARS-CoV-2 B.1.617.2 Spike GCN4-IZ Protein, CF (catalog number 10878-CV).

### Enzyme-linked immunospot (ELISpot) assay

Cellular immunity was assessed by a standard ELISpot assay using a pool of peptides derived from a peptide scan for each protein used as an antigen. For ELISpot assay, mice were sacrificed and spleens collected. Spleens were homogenized in HBSS and washed with 10% CTL Wash-RPMI medium and erythrocytes were lysed by BD Pharm Lyse (BD Biosciences). Next, a mouse IFN-γ/IL-4 double-color ELISPOT kit (Cellular Technology Limited, CTL, Ohio, USA) was used according to manufacturer’s instructions. Briefly, splenocytes (2 × 10^5^/well) were incubated in CTL-test medium (with 1% GlutaMax) with or without 1μg/ml of pooled peptides for a specific antigen. After overnight incubation, IFN-γ/IL-4 secreting splenocytes were detected using anti-murine IFN-γ (FITC) or IL-4 (Biotin) antibodies, respectively. Spots were developed and counted with an S6 entry M2 ELISPOT reader. For each vaccine antigen, the following peptide libraries were used:

- 15mer peptides for RBD of SARS-CoV-2 (original strain): a pool of 53 peptides derived from a peptide scan (15mers with 11 amino acid overlap) through the RBD of SARS-CoV-2 (an original Wuhan strain) [JPT Peptides: PepMix SARS-CoV-2 (S-RBD) PM-WCPV-S-RBD-2].
- 9mer peptides for RBD of SARS-CoV-2 (original strain): a pool of 215 peptides derived from a peptide scan (9mers with 8 amino acid overlaps) through the RBD of SARS-CoV-2 (an original Wuhan strain). The peptides were custom-made by JPT Peptides.
- 15mer peptides for RBD of SARS-CoV-2 (omicron variant): a pool 53 peptides derived from a peptide scan (Peptide scan (15mers with 11 amino acid overlap) through RBD of Spike Glycoprotein - contains mutations G0339D, S0371L, S0373P, S0375F, K0417N, N0440K, G0446S, S0477N, T0478K, E0484A, Q0493R, G0496S, Q0498R, N0501Y, Y0505H of SARS-CoV-2. [JPT Peptides: PepMix PM-SARS2-RBDMUT08-1]
- 15mer peptides for spike of MERS-CoV: a pool of 336 (168+168) peptides derived from a peptide scan (15mers with 11 amino acid overlap) through the spike glycoprotein (Swiss-Prot ID: K9N5Q8) of MERS-CoV (Middle East respiratory syndrome-related coronavirus) [JPT peptides Product Code: PM-MERS-CoV-S-1].
- 15mer peptides for nucleoprotein of SARS-CoV-2 (original strain): a pool of 102 peptides derived from a peptide scan (15mers with 11 amino acid overlap) through nucleoprotein (UniProt: P0DTC9) of SARS-CoV-2 [JPT peptide Product Code: PM-WCPV-NCAP].
- 15mer peptides for nucleoprotein of MERS-CoV: a pool of 101 peptides derived from a peptide scan (15mers with 11 amino acid overlaps) through the nucleoprotein of MERS-CoV (YP_009047211.1). The peptides were custom-made by JPT Peptides.

### Intracellular cytokine staining and flow cytometry analysis

Analyses were performed using the following antibodies: CD3-FITC (145-2C11, Biolegend), CD4-BV750 (GK1.5, Biolegend), CD8-BV786 (53-6.7, Biolegend), CD62L-BV510 (MEL-14, Biolegend), CD44-PE/Cy5 (IM7, Biolegend), CD107a-BV711 (1D4B, Biolegend), CD154-PE (MR1, BD), Gmzb-AF700 (QA16A02, Biolegend), IFN-γ-APC (XMG1.2, Biolegend) and TNF-α-BB700 (MP6-XT22, BD). In brief, splenocytes isolated from experimental mice were cultured in the presence of phorbol 12-myristate 13-acetate (PMA) (1.9 nM) and ionomycin (0.08 mg/ml) solution at 37°C for 4 hours. After first washing, 2 × 10^6^ cells were stained with FVS780 (viability dye) and followed by antibodies for surface antigens (Anti-CD3, CD4, CD8, CD62L, CD44, CD107a, and CD154) in 4°C for 30 minutes. After washing, cells were fixed and permeabilized with BD Cytofix/Cytoperm, and then stained with the antibodies against the intracellular antigens (Gzmb, IFN-γ, and TNF-α) for 20 minutes at 22°C. Stained cells were separated using a BD FACSymphony™ Cell Analyzer and one million events were collected for each sample. The analysis was performed in FlowJo software version 10.6.2.

### ELISA assay

Submandibular blood was collected for ELISA assay. Serum IgG against RBD of SARS-CoV-2 (original strain) was measured by ELISA assay using ENZO SARS-CoV-2 IgG ELISA Kit [Cat# ENZ-Kit109-0001]. Serum IgG against nucleoprotein of SARS-CoV-2 (original strain) was measured by ELISA assay using ENZO ENZ-KIT193-0001. An ELISA kit was purchased from ENZO Lifesciences, 100 ng of Mouse COVID-19 spike RBD Domain Coronavirus Monoclonal Antibody from MyBiosource (Cat# MBS434247) was used as a positive control, while serum from unvaccinated mice was pooled and used as a negative control. Assays were performed according to manufacturer’s instructions. In brief, mouse serum was diluted 1:100 in sample diluent and anti-RBD antibody was diluted to 1.0ug/ml with sample diluent. A 100ul aliquot was added to each designated well and incubated at 37°C for 30min. Post incubation, the plate was washed 4 times with 1X wash buffer using an automatic washing machine (Bio Tek TS-50), Then anti-Mouse IgG Fc Secondary antibody (Invitrogen Cat#31437) was diluted in HRP dilution buffer (Sigma Cat#18995) to 1:5000 and 100ul added to each well. The plate was then incubated at 37°C for 15 min. Following incubation, the plate was washed 4 times with 1X wash buffer. Subsequently, 100ul of 3,3′,5,5′-Tetramethylbenzidine (TMB) substrate solution was added to each well and the plate incubated in the dark at 37°C for 15 min. The reaction was stopped by addition of 50ul of stop solution per well and absorbance values were measured at 450 nm using a plate reader (TECAN, Spectrafluor plus).

### PRNT_50_ assay

For the PRNT assay, Vero76 cells were first treated with serially diluted mouse serum, followed by infection with live virus of SARS-CoV-2 (Delta variant strain). In this assay, the infected cells die and form a plaque after fixation and staining with crystal violet. If the serum contains neutralizing antibodies, the viral infection is inhibited, resulting in a reduction of the number of plaques. The results are shown as the dilution titer of serum that show 50% reduction of number of plaques (PRNT_50_). The assay was performed by BIOQUAL, Inc. (Rockville, MD).

### In vivo cell elimination assay

4T1 (ATCC, CRL-2539) and 4T1-Luc2 (ATCC, CRL-2539-Luc2) mouse mammary tumor cell lines, derived from BALB/c mice, were cultured in RPMI medium supplemented with 10% FBS at 37°C in a humidified atmosphere containing 5% CO2. For 4T1-Luc2 cells, the medium was supplemented with 8 μg/ml blasticidin (Invivogen). 4T1 and 4T1-Luc2 cells were transfected with a plasmid DNA encoding a fusion protein of the nucleoproteins of SARS-CoV-2 and MERS-CoV (non-secreted form, i.e., without CD5 signal peptide) under the CMV promoter and the hygromycin-resistant gene under early SV40 promoter control, using lipofectamine 2000. Cells expressing the fusion protein of the nucleoproteins of SARS-CoV-2 and MERS-CoV (called 4T1-SMN), were isolated by culturing the cells in the presence of 200 μg/mL of hygromycin B. Transfected cells were selected with hygromycin at a final concentration of 50~200μg/ml and several colonies were picked and expanded for further analysis. 1×10^5^ 4T1-Luc2 (spike) and 5×10^4^ 4T1 (SMN) cells in 50μl PBS were injected into the 4^th^ mammary fat pad of BALB/c under isoflurane anesthesia and mice were observed daily. Tumor volume (length × width^2^ × 0.5) was measured by digital caliper and recorded.

