## Supplemental Figures for "Controllable self-replicating RNA vaccine delivered intradermally elicits predominantly cellular immunity"

### Title

### Affiliations

Minoru S. H. Ko, MD, PhD

Elixirgen Therapeutics, Inc.

Science + Technology Park at Johns Hopkins

855 N. Wolfe Street, Suite 624

Baltimore, MD 21205, USA

**FIG. S1. In vitro expression of c-srRNA1-GFP at 33°C and at 37°C.**

mRNAs, either synthetic modified mRNA encoding GFP (mRNA-GFP [5mC, ψ]), T7-VEE-GFP-ts, or c-srRNA1-GFP, were tested for their GFP expression using human adult dermal fibroblast cells (HDFa). 50,000 cells in each well of 24-well plate were transfected with 0.5 µg of mRNA with Lipofectamine MessengerMAX (Day 0) and cultured in the DMEM medium containing 500 ng/ml B18R at 33°C or 37°C.

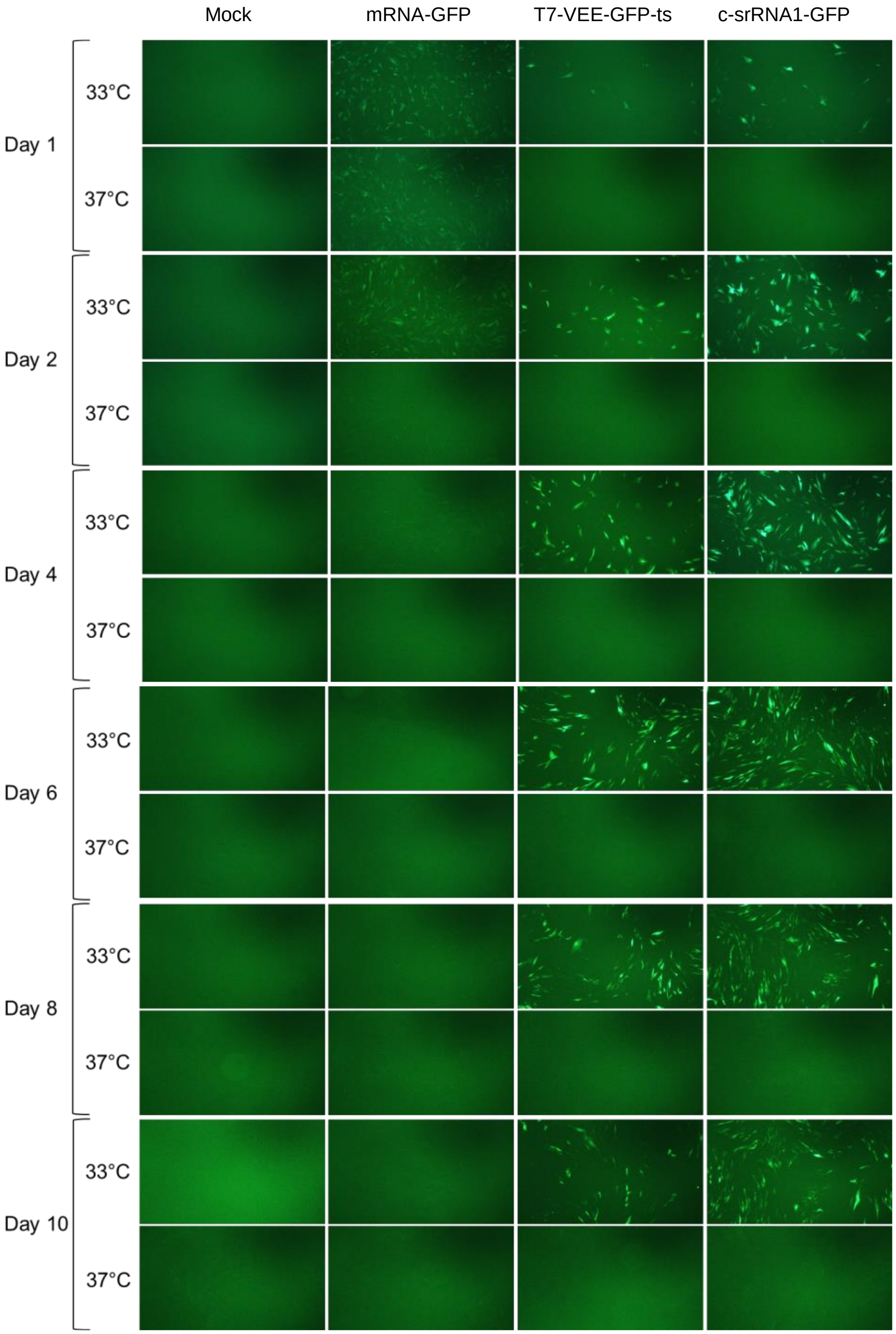

**FIG. S2. c-srRNA functions in skin by intradermal injection, but not in muscle by intramuscular injection.**

**(A)** 5 µg of c-srRNA3-LUC as naked RNAs without LNP was injected intradermally onto the shaved skin or intramuscularly into BALB/c mice (Day 0). Luciferase activity was visualized and quantitated by using a bioluminescent imaging system from Day 1 through Day 9 post-injection. Images of Day 1 and Day 9 are shown. (**B**) Luciferase activities are plotted from Day 1 through Day 9. Left: Results from intradermal injection. Right: Results from intramuscular injection.

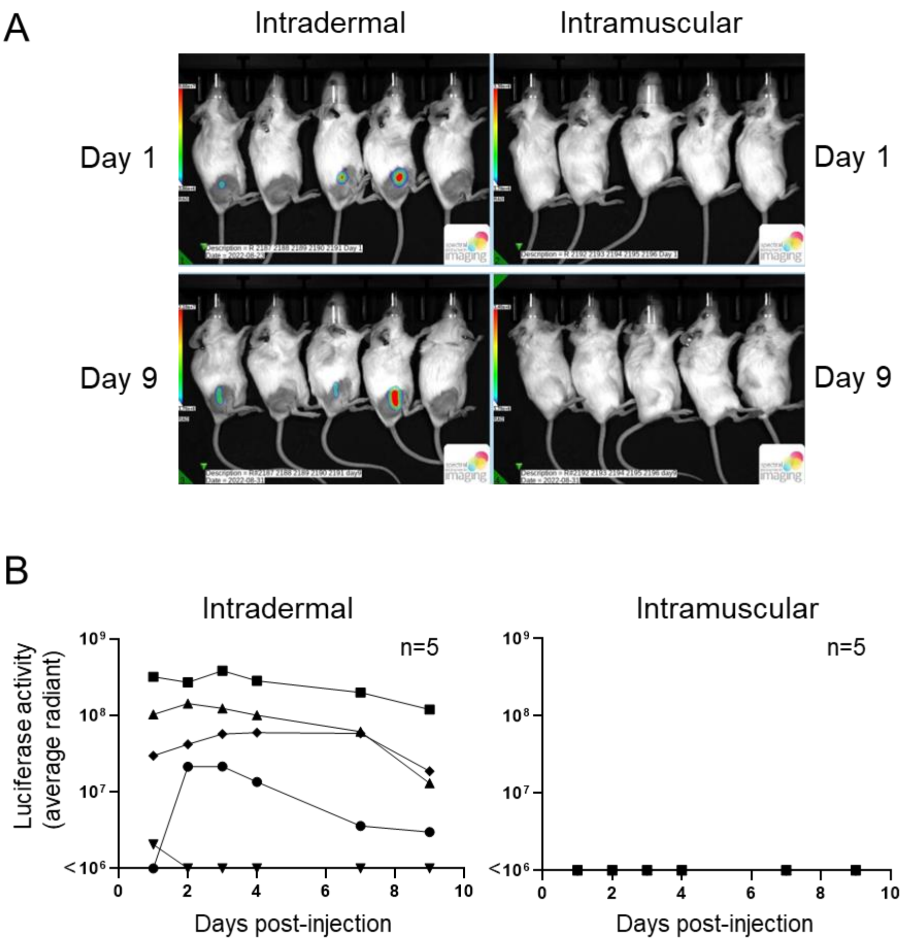

**FIG. S3. Comparison of cellular immunity induced by EXG-5006a and EXG-5006b.**

**(A)** A schematic diagram of experimental procedures. BALB/c mice were immunized by a single intradermal injection of 100 µL solution containing either a placebo (PBO: buffer only: n=4 or 5), 5 µg (n=5), 25 µg (n=5) of EXG-5006a (non-codon optimized) or EXG-5006b (codon-optimized). After 14 days, mice were sacrificed and splenocytes were collected for ELISpot assays. (**B**) Results of ELISpot assays performed by restimulating splenocytes in the presence or absence of a pool of SARS-CoV-2 nucleoprotein peptides. The frequency of IFN-γ- or IL-4-secreting cells obtained in the presence of peptides is plotted in the graph after subtracting the frequency obtained in the absence of peptides (background). The average and standard deviation (error bars) are shown for each group.

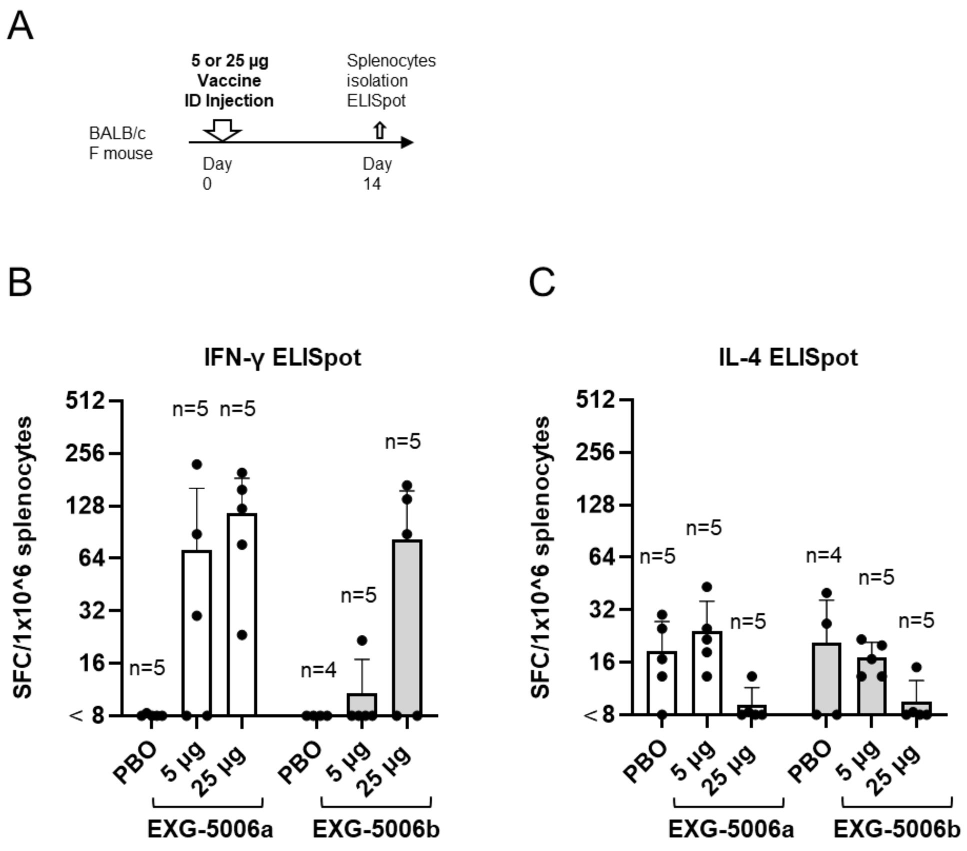
